# Title: *In vivo* phosphatidylserine variations steer Rho GTPase signaling in a cell-context dependent manner

**DOI:** 10.1101/471573

**Authors:** Matthieu Pierre Platre, Vincent Bayle, Laia Armengot, Joseph Bareille, Maria Mar Marques-Bueno, Audrey Creff, Lilly Maneta-Peyret, Jean-Bernard Fiche, Marcelo Nolmann, Christine Miège, Patrick Moreau, Alexandre Martinière, Yvon Jaillais

## Abstract

**Abstract:** Rho GTPases are master regulators of cell signaling, but how they are regulated depending on the cellular context is unclear. Here, we show that the phospholipid phosphatidylserine acts as a developmentally-controlled lipid rheostat that tunes Rho GTPase signaling in Arabidopsis. Live super-resolution single molecule imaging revealed that RHO-OF-PLANT6 (ROP6) is stabilized by phosphatidylserine into plasma membrane (PM) nanodomains, which is required for auxin signaling. Furthermore, we uncovered that the PM phosphatidylserine content varies during plant root development and that the level of phosphatidylserine modulates the quantity of ROP6 nanoclusters induced by auxin and hence downstream signaling, including regulation of endocytosis and gravitropism. Our work reveals that variations in phosphatidylserine levels are a physiological process that may be leveraged to regulate small GTPase signaling during development.

**One Sentence Summary:** Phosphatidylserine acts as a developmentally-controlled lipid rheostat that regulates cellular auxin sensitivity and plant development.

## Main Text

Proteins from the Rho/Ras superfamily are small GTPases that regulate fundamental eukaryotic functions, including cell signaling, cell polarity, intracellular trafficking and cytoskeleton dynamics (1, 2). Furthermore, they control the morphology and behavior of cells and organisms by integrating signaling pathways at the cell surface into various cellular outputs. The small GTPase paradigm stipulates that they are in an “inactive” form when bound to GDP, and in an “active” form when bound to GTP. However, emerging evidence suggest that this view is likely oversimplified, since their membrane environment also dictates the signaling capacity of these GTPases (2, 3). In particular, Ras/Rho signaling is intimately linked with membrane lipids in all eukaryotes. Interaction with anionic lipids is important for their plasma membrane (PM) targeting (4, 5), but also mediates the clustering of these small GTPases at the cell surface into nanometer scale membrane domains (6-8). Phosphoinositides are low abundant anionic phospholipids that can be acutely produced or metabolized by dedicated enzymes with exquisite subcellular precision, and as such often function as signaling lipids (9). Moreover, they mediate the recruitment of some Ras/Rho proteins to the cell surface and into nanoclusters (4, 7, 10). Phosphatidylserine (PS) is also involved in the nanoclustering and signaling of some GTPase, such as K-Ras in human and Cdc42 in yeast (3, 6, 8, 11-13). However, by contrast to phosphoinositides, PS is a relatively abundant anionic phospholipid, representing up to 10-20% of the total phospholipids at the PM inner leaflet (14). In addition, PS is not constantly modified by specialized metabolizing enzymes and the subcellular PS repartition is thought to be relatively stable across cell types (14). Therefore, PS appears to be a structural component of the membrane, which is required for K-Ras/Cdc42 nanoclustering. It is unknown, however, whether PS also has a regulatory role *in vivo* in modulating nanocluster formation and subsequent signaling. In other words, is PS function in GTPase nanoclustering rate limiting? And if yes, are PS levels regulated during development and what are the consequences of such changes on small GTPases signaling capacity? Here, we addressed these questions using the *Arabidopsis thaliana* root as a model system, because it is a genetically tractable multicellular organ, with a variety of cell types and cell differentiation states and amenable to live imaging, including super-resolution microscopy (15).

In plants, there is a single protein family in the Ras/Rho GTPase superclade, called ROP for RHO-OF-PLANT(16). ROPs are master regulators of cell polarity and cell morphogenesis, but they also sit at the nexus of plant hormone signaling (including auxin and abscisic acid), cell wall sensing pathways and receptor-like kinase signaling (involved in development, reproduction and immunity) (16-27). Here, we show that auxin triggers ROP6 nanoclustering within minutes, in a PS dependent manner. Furthermore, we found that PS is required for ROP6 signaling, and variations in the cellular PS content directly impact the quantity of ROP6 nanoclusters and thereby subsequent downstream auxin signaling, including the regulation of endocytosis and root gravitropism. Therefore, PS is not a mere structural component of the membrane, it is a *bona fide* signaling lipid that acts as a developmentally-controlled lipid rheostat to regulate small GTPases in a cell-context dependent manner

## Results and discussion

### Plasma membrane phosphatidylserine levels vary during root cell differentiation

Phosphatidylserine (PS) is an anionic phospholipid that partitions between the cytosolic leaflets of the PM and endosomes (28). Bulk PS measurement in *Arabidopsis thaliana* suggested that the relative PS concentration can vary *in vivo* depending on the organ (29). In order to get tissue and cellular resolution on the PS distribution, we recently validated the use of two PS reporters in Arabidopsis (28, 30), the PS-binding C2 domain of Lactadherin (C2^Lact^) (5) and the PH domain of EVECTIN2 (2xPH^EVCT2^) (31). In both cases, the proportion of PS sensors was markedly more pronounced at the PM than endosomes in the root basal meristem compared to cells in the elongation zone (Fig. 1A, Fig. S1A-B). This developmental gradient appeared to be in part regulated by the plant hormone auxin as relatively short treatment (60min) with the synthetic auxin naphthalene-1-acetic acid (NAA) increased the level of both PS sensors at the PM at the expense of their endosomal localization in the elongation zone (Fig. 1B, Fig. S1C). Therefore, not only the overall PS level vary depending on the organ but there are also local variations of the PS content at the PM within an organ, during cell differentiation and in response to hormonal cues.

**Figure 1.**
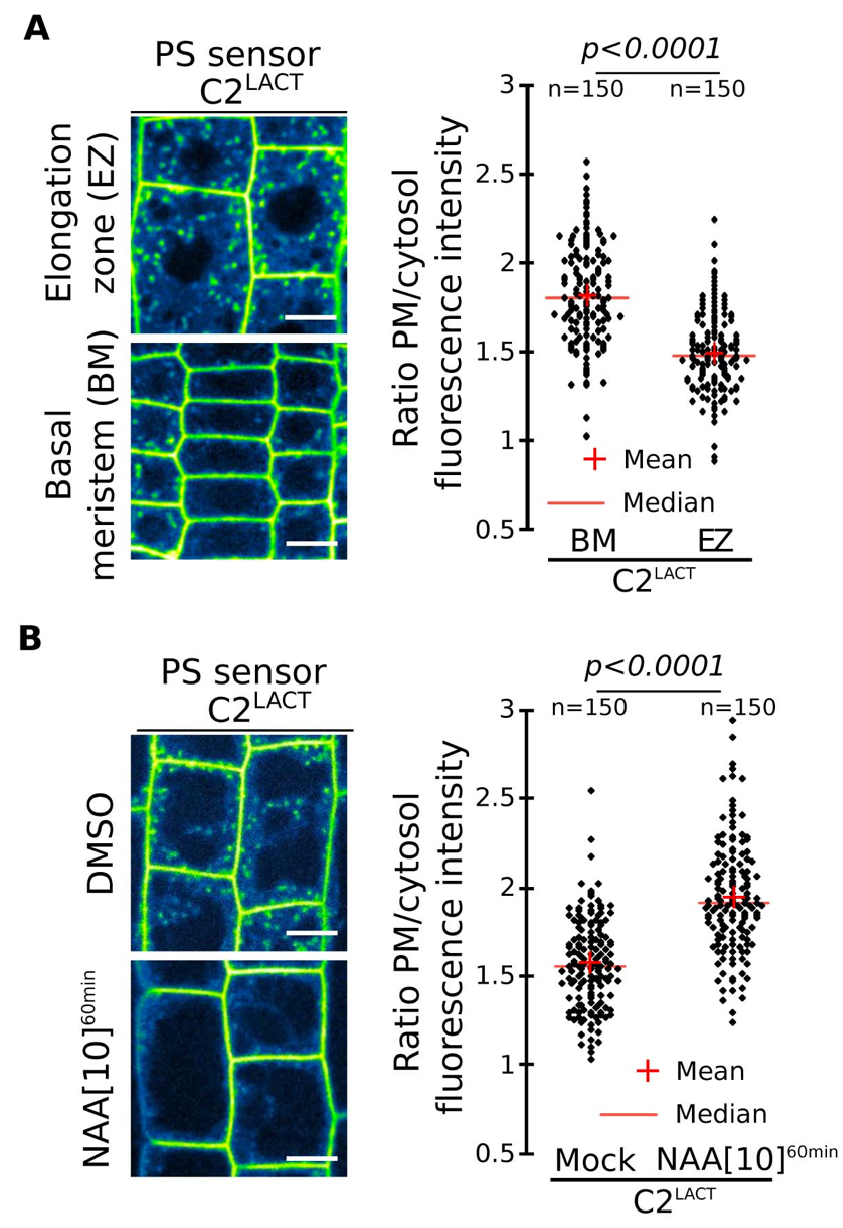
Plasma membrane phosphatidylserine levels vary during root cell differentiation. A, Confocal images of mCIT-C2^LACT^ root epidermis in basal meristem (BM) and elongation zone (EZ) and related quantification. Confocal images of mCIT-C2^LACT^ root epidermis in the absence/presence of NAA (10μM, 60min) and related quantification. Scale bars 10μm. n represents the number of cells, letters indicate statistical differences (see methods for details on statistical tests).

### Graded phosphatidylserine levels tune ROP6 signaling

In order to test the potential impact of PS variations during development, we experimentally manipulated the plant PS content, from no PS biosynthesis in the *phosphatidylserine synthase1* (*pss1*) mutant (28), to mild PS levels in transgenic lines expressing artificial microRNAs against *PSS1* (*amiPSS1*), and high PS levels in transgenic lines overexpressing *PSS1* (*PSS1-OX*) (Fig. S1D-E). The changes in PS content measured in *amiPSS1* and *PSS1-OX* lines of about ±2-fold fell well into the physiological range, since PS levels in Arabidopsis vary about 5-fold between roots and leaves tissues (29). The *pss1* mutant showed defects in root gravitropism (Fig. S1F-G). Quantitative analyses of root bending following gravistimulation (Fig. S1H) revealed that the *pss1-3* mutant had no gravitropic response (Fig. 2A), *amiPSS1* lines had an attenuated response, while *PSS1-OX* lines were hypergravitropic (Fig. 2B). These opposite gravitropic phenotypes of *PSS1* loss- and gain-of-function resembled those of ROP6, a Rho-Of-Plant (ROP) GTPase, which is activated by auxin and regulates root gravitropism (*21, 22*). Like *PSS1-OX* lines, lines overexpressing either ROP6 (*ROP6-OX*) or constitutive active GTP-lock ROP6 (*ROP6^CA^*) showed a hypergravitropic phenotypes, which were abolished in a *pss1-3* background (Fig 2A). During root gravitropism, ROP6 acts downstream of auxin to inhibit endocytosis and regulate microtubule orientation (21, 22, 32, 33). Similar to *rop6 (21, 22, 32*), we observed that in *pss1-3*: i) FM4-64 and PIN2-GFP uptake in the presence of BrefeldinA (BFA) was increased (Fig. 2C, Fig. S2A-D), ii) auxin failed to inhibit FM4-64 and PIN2-GFP endocytosis (Fig. 2C, Fig. S2A-D), iii) CLATHRIN-LIGHT-CHAIN2 (CLC2)-GFP PM association was insensitive to auxin treatment (Fig. S2E), and iv) auxin-triggered microtubule reorientation was abolished (Fig. S2F). FM4-64 uptake in *pss1-3xROP6^CA^* was identical to that of *pss1-3* single mutant and opposite to *ROP6^CA^*(Fig. 2C), showing that PSS1 is required for ROP6-mediated inhibition of endocytosis. Furthermore, transgenic lines with low PS content (*amiPSS1*) had decreased auxin-mediated inhibition of endocytosis, while lines with heightened-PS content (*PSS1-OX*) mimicked *ROP6^CA^* phenotypes with pronounced inhibition of endocytosis upon auxin treatment (Fig. 2D, Fig. S2G-I). Together, our analyses suggest that i) PS is required for auxin-mediated ROP6 signaling during root gravitropism and ii) PS levels impact the strength of ROP6 signaling output in a dosedependent manner.

**Figure 2.**
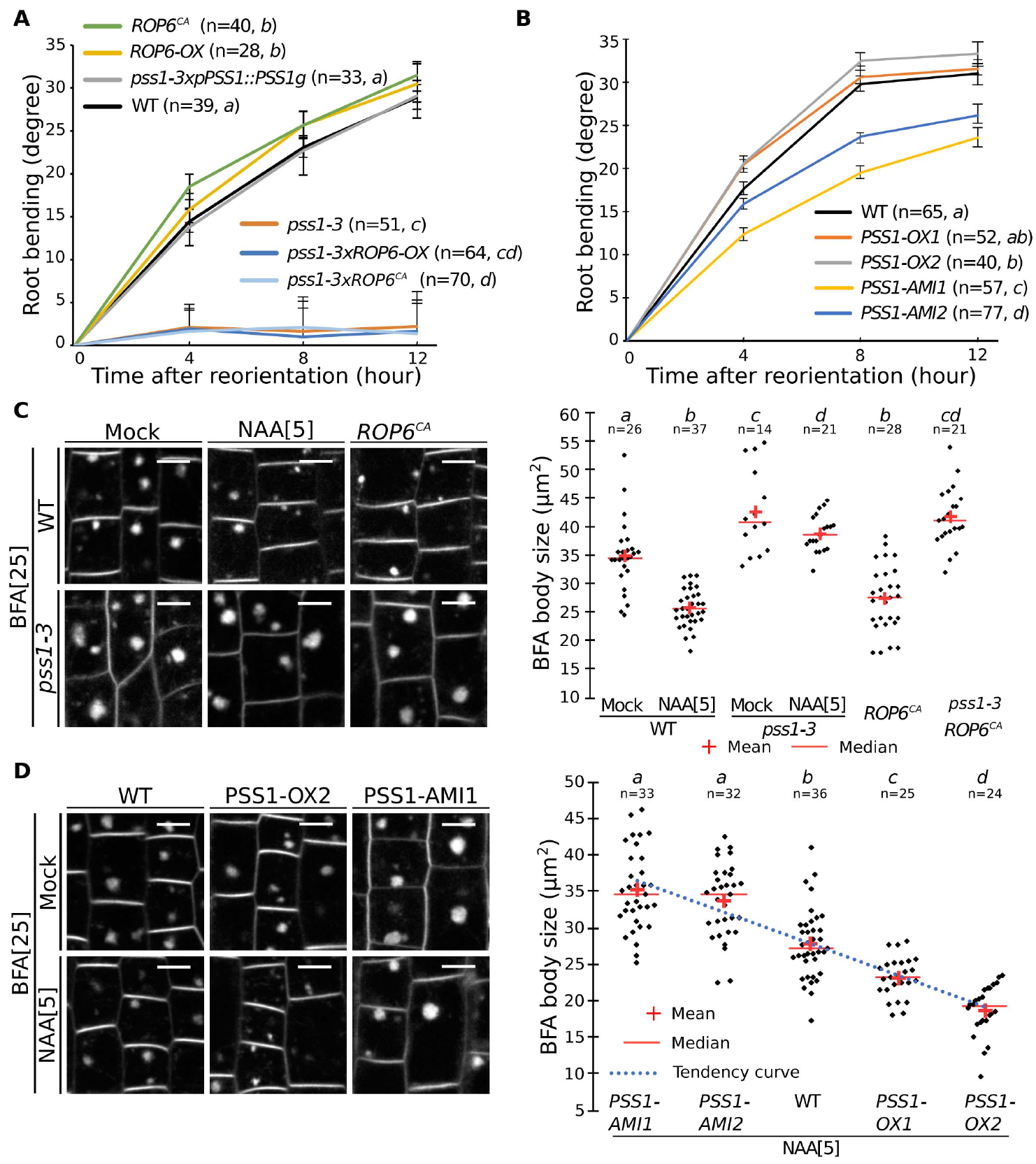
Variation in phosphatidylserine concentration tunes ROP6 signalling. **A-B**, Quantification of root bending after gravistimulation (mean +/- sem). C-D, Confocal images of FM4-64 staining in root epidermis (BFA: 25μM; NAA: 5μM), and related quantification. Scale bars 10μm. n represents the number of roots analysed, letters indicate statistical differences (see methods for details on statistical tests).

### Auxin triggers ROP6 nanoclustering at the plasma membrane

PS and ROP6 both accumulate at the PM, we therefore reasoned that PS may contribute to ROP6 localization. Surprisingly, GFP-ROP6 localization, as seen by confocal microscopy, was almost identical in *pss1-3* and WT, being mainly at the PM and only faintly delocalized in intracellular compartments in *pss1-3* (Fig. S3A). In leaves, ROP6^CA^ was previously shown to be confined in membrane domains (34), raising the possibility that PS could contribute to ROP6 signaling by regulating its lateral segregation at the PM. To analyze ROP6 PM partitioning in root cells and in the context of auxin response, we developed several microscopy-based assays, including Fluorescence Recovery After Photobleaching (FRAP), Total Internal Reflection Fluorescence Microscopy (TIRFM) (35) and PhotoActivated Localization Microscopy (PALM) (15) (Fig. S4). As shown for ROP6^CA^ in leaf (34), activation of ROP6 (here using auxin treatment) delayed GFP-ROP6 fluorescence recovery after photobleaching (Fig. 3A-B and Fig. S3B-D). TIRFM on root tip epidermal cells allowed to focus only on the plane of the PM with a 100nm axial resolution (35) (Fig. S4B) and revealed that GFP-ROP6 mostly localized uniformly at the PM (Fig. 3C). By contrast, in auxin-treated plants, GFP-ROP6 additionally resided in diffraction-limited spots present in the plane of the PM (Fig. 3C), suggesting that auxin treatment triggers the clustering of ROP6 in membrane domains. By using stochastic photoswitching on live roots, single particle tracking PALM (sptPALM) experiments provided tracks of single molecule localization through time, and therefore allowed us to analyze the diffusion behavior of single ROP6 molecule in response to auxin (Fig. S4D). While mEos-ROP6 molecules in the untreated condition were almost exclusively diffusing, mEos-ROP6 molecules in plants treated for 5 minutes with auxin (or mEos-ROP6^CA^ molecules) existed in two states at the PM of epidermal cells: immobile or diffusing (Fig. 3D-E, Fig. S5A-C and Supplementary Video 1). Clustering analyses on live PALM images (36, 37) showed that auxin triggered the clustering of mEos-ROP6 in PM-nanodomains of about 50 to 70 nm wide (Fig. 3F-G and Fig. S6). Together, our data indicate that activation, either genetically (i.e. ROP6^CA^) or by an endogenous activator (i.e. auxin), triggers ROP6 recruitment, immobilization and stabilization into PM-nanodomains and that these events happen minutes following auxin treatment.

**Figure 3.**
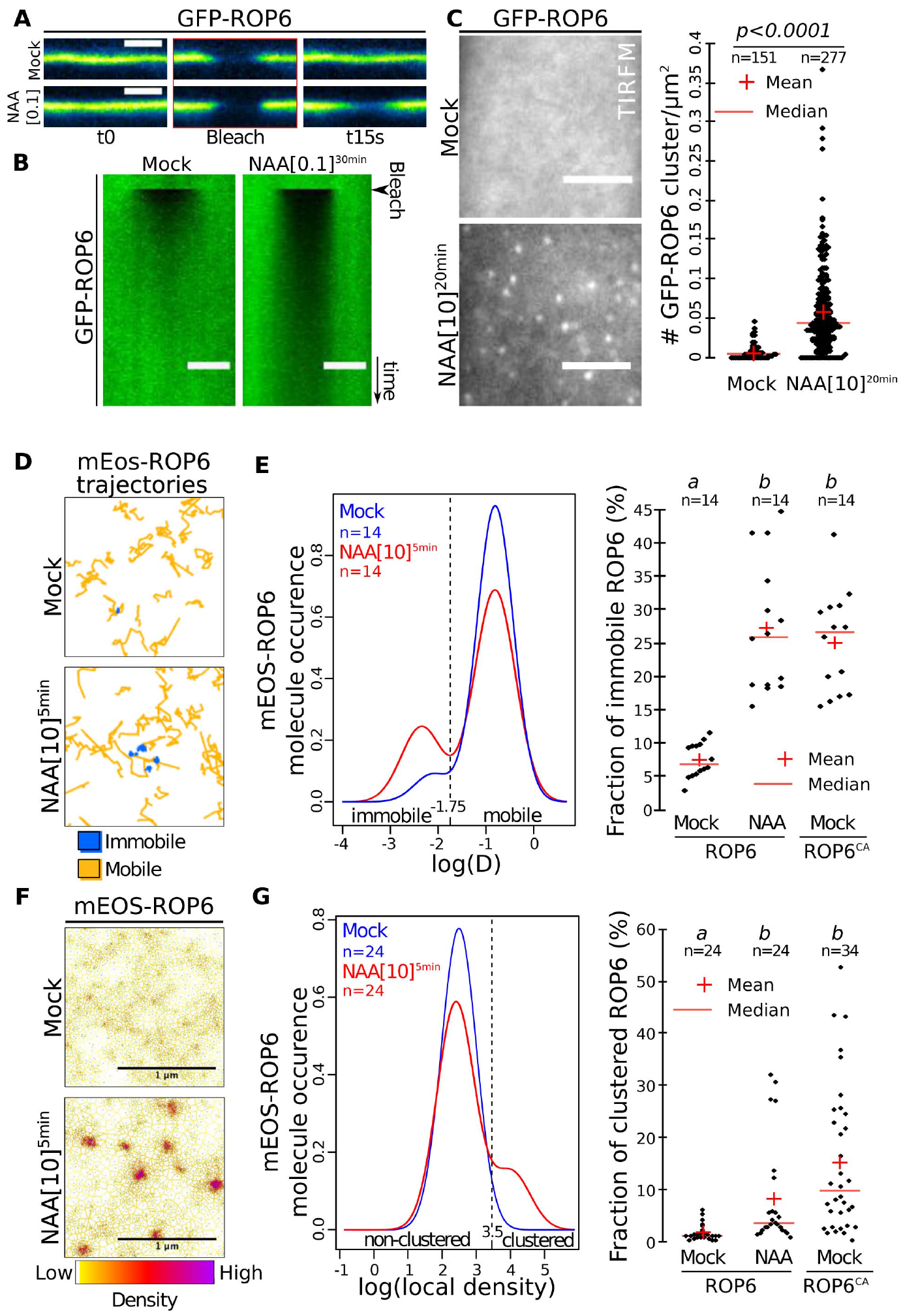
Auxin triggers ROP6 nanoclustering at the PM of root epidermal cells. A, Confocal images of GFP-ROP6 during FRAP experiments (NAA: 100nM, 30 min), **B**, related kymograph (time scale 9 seconds). **C**, TIRFM micrograph of GFP-ROP6 (NAA: 10μM, 20 min) and related quantification. **D**, Representative mEos-ROP6 trajectories obtained by sptPALM analyses.**E**, Distribution of mEos-ROP6 molecules according to their apparent diffusion coefficient (D) obtained by analysing sptPALM trajectories (NAA: 10μM, 5 min), and related quantification. **F**, Live PALM analysis of mEos-ROP6 molecules density by tessellation-based automatic segmentation of super-resolution images (NAA: 10μM, 5 min). **G**, Distribution of mEos-ROP6 molecules according to their local density and related quantification. Scale bars: 5 μm (A-C), 1 μm (F). n represents the number of ROIs (C), or independent acquisitions (different cells) (E and G) analysed, letters indicate statistical difference (see methods for details on statistical tests).

### PS regulates auxin-induced ROP6 nanoclustering in a dose-dependent manner

Next, we tested the impact of PS on ROP6 PM dynamics. In FRAP experiments, GFP-ROP6 sensitivity to auxin was reduced in *pss1-3* (Fig. 4A and Fig. S3B-E), suggesting that PS is critical for the immobilization of ROP6 by auxin. In WT plants, NAA-induced GFP-ROP6 presence in PM-nanodomains was more pronounced in the basal meristem than in the elongation zone in TIRFM experiments (Fig. 4B), which correlated with the observed differential presence of PS content at the PM in these regions (Fig. 1A). To analyze whether this differential auxin sensitivity was dependent on the amount of PS present in these cells, we performed PS loss- and gain-of-function experiments. First, auxin failed to induce GFP-ROP6 nanodomains in both region of the root in *pss1-3* (Fig. 4C), suggesting that PS is indeed required for auxin-triggered ROP6 nanoclustering. Second, exogenous treatment with lysophosphatidylserine (lyso-PS), a more soluble lipid than PS but with an identical head group (28), boosted the number of auxin-induced GFP-ROP6 nanodomains observed in TIRFM in WT plants (Fig. 4D). Together these data suggest that the quantity of PS at the PM impacts ROP6 nanoclustering. While PS was required for auxin-triggered ROP6 nanoclustering, a certain amount of ROP6 was still found in PM domains in *pss1*, independent of the presence of auxin (Fig. 4C). Kymograph analyses revealed that ROP6-containing PM-nanodomains observed by TIRFM were immobile in both WT and *pss1-3* (Fig. 4E). Photobleaching experiments showed that GFP-ROP6 was highly stable in these PM-nanodomains in the WT (i.e. no fluorescence recovery of GFP-ROP6 in PM-nanodomains, by contrast to a fast recovery of fluorescence outside of these domains) (Fig. 4E and Supplementary Video 2). By contrast, GFP-ROP6 fluorescence in PM-nanodomains was rapidly recovered in *pss1-3*, suggesting that ROP6 was not stabilized into PM-nanodomains in the absence of PS (Fig. 4E and Supplementary Video 3). Together, our results show that PS is necessary for both ROP6 stabilization into PM-nanodomains and downstream ROP6 signaling, including regulation of endocytosis and root gravitropism.

**Figure 4.**
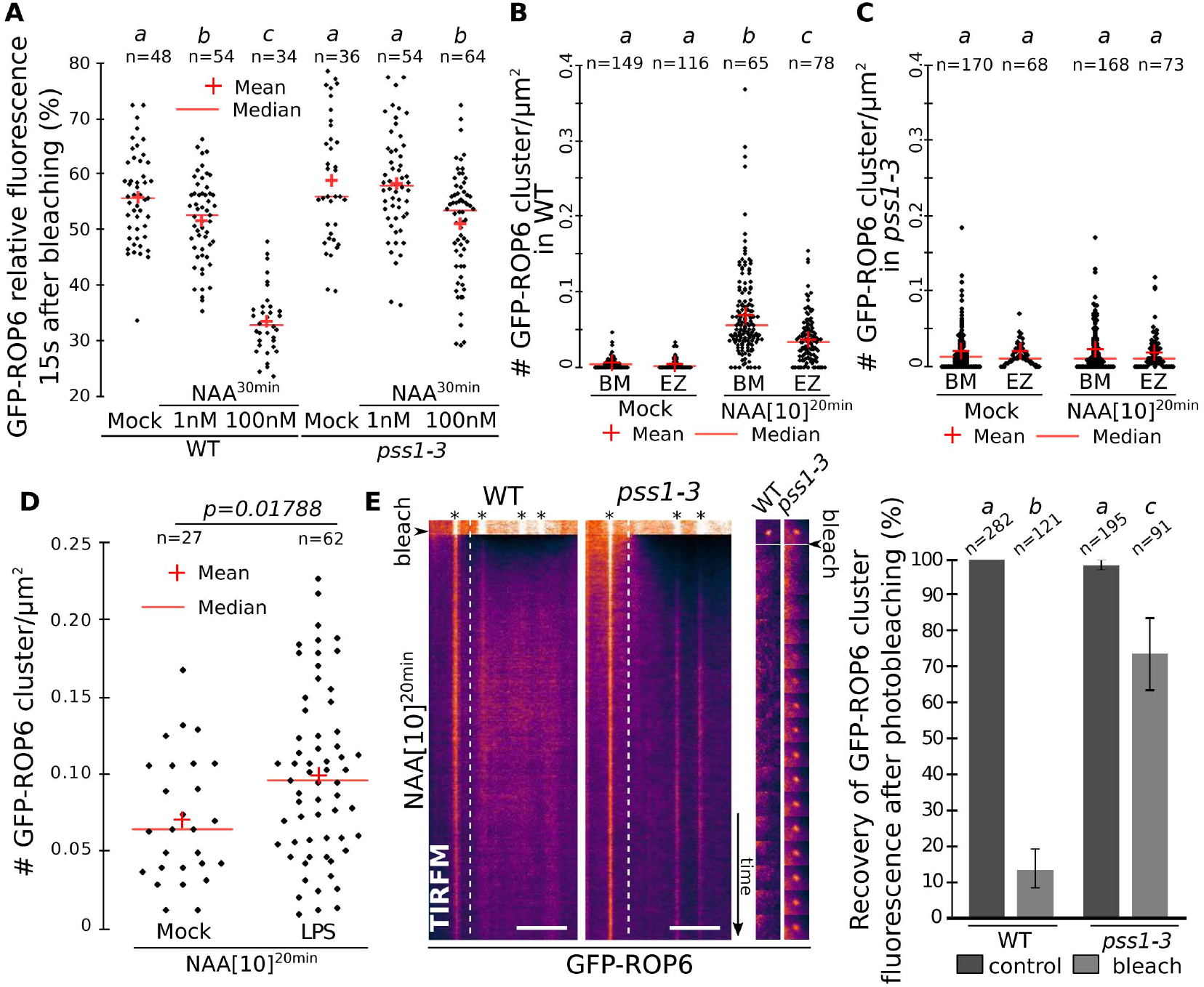
Phosphatidyserine is necessary for auxin-induced stabilization of ROP6 into PM-nanodomains. A, Quantification of FRAP experiments in WT and *pss1-3* root epidermal cells (NAA: 1nM and 100nM, 30 min). **B-C**, Quantification of TIRFM experiment in WT and *pss1-3* root epidermal cells (NAA, 10μM, 20 min) in basal meristem (BM) and elongation zone (EZ) (data are the same as in Fig3C but split into its respective zone). **D**, Quantification of TIRFM experiment in the presence or absence of lyso-PS in the meristematic region (NAA: 10μM, 20min). **E**, Kymograph of GFP-ROP6 localization obtained by TIRFM, (time scale 12 seconds) with images of a single GFP-ROP6 nanocluster (9s interval) and related quantification. Scale bars: 5 μm. n represents the number of ROIs (A-D), or GFP-ROP6 nanodomains (E) analysed, letters indicates statistical difference (see methods for details on statistical tests).

### Immobile phosphatidylserine molecules accumulate in PM-nanodomains together with ROP6

Next, we addressed whether the regulation of ROP6 clustering and signaling by PS was direct. If so, we should expect PS, like ROP6, to also localize in PM-nanodomains. Using sptPALM and clustering analyses, we found that i) the PS reporter mEos-2xPH^EVCT2^ segregated into nanodomains at the PM of root epidermal cells (Fig. 5A) and ii) about 35% of mEos-2xPH^EVCT2^ molecules were present as a slow-diffusible population (Fig. 5B and Fig S5D), showing an apparent diffusion coefficient similar to that of immobile mEos-ROP6 (Fig. 3E) and suggesting that PS and ROP6 may co-exist in the same PM-nanodomains. Accordingly, co-visualization of GFP-ROP6 and the PS sensor 2xmCHERRY-C2^Lact^ in TIRFM confirmed that they at least partially reside in the same PM-nanodomains in response to auxin (Fig. 5C).

**Figure 5.**
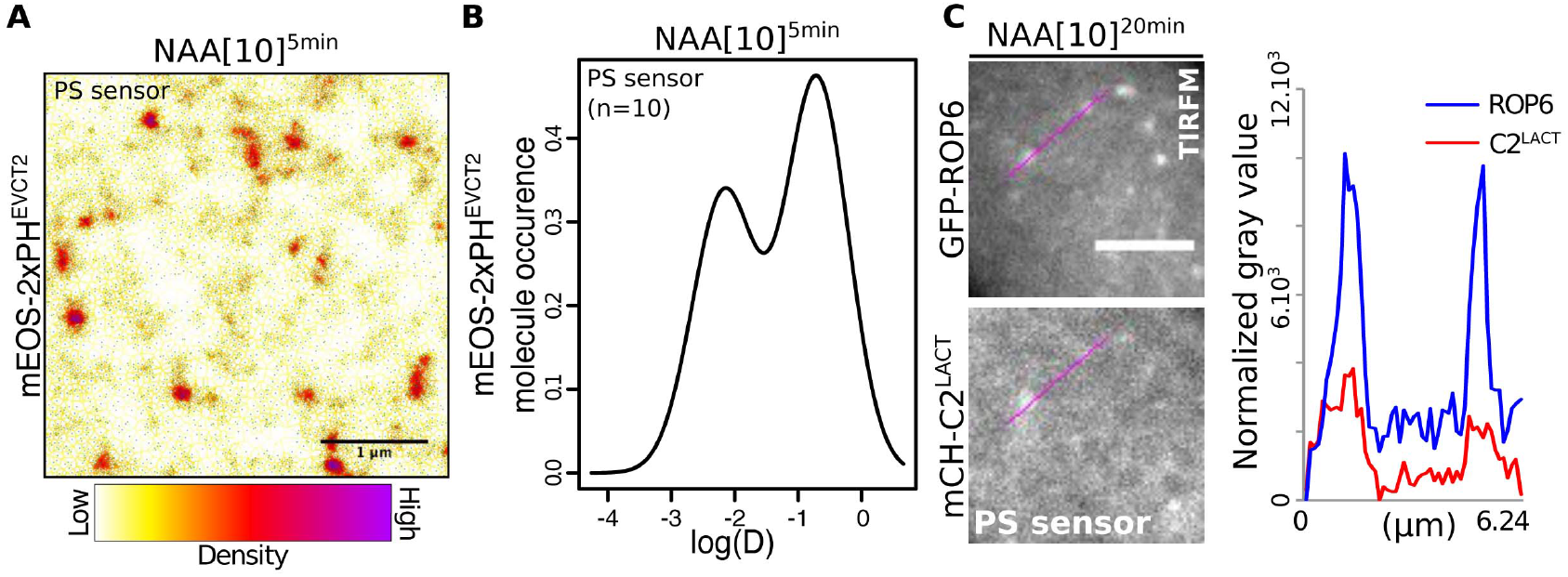
Phosphatidylserine concentrates in nanoclusters at the PM. A, live PALM analysis of mEos-2xPH^EVCT2^ localization (NAA, 10μM, 5 min) in root epidermal cells and **B**, distribution of mEos-2xPH^EVCT2^ molecules according to their diffusion. **C**, TIRFM micrograph of root cells co-expressing GFP-ROP6 with the PS marker 2xmCH-C2^LACT^ (NAA: 10μM, 20 min) and line scan of fluorescence intensities. Scale bars 1μm (A), 5μm (C). n represents the number of acquisitions (independent cells) (B), letter indicates statistical difference (see methods for details on statistical tests).

### ROP6 interaction with anionic phospholipids is required for nanoclustering and downstream signaling

ROP6 possess in its C-terminus a polybasic region adjacent to a prenylation site (Fig. S7A). Such polybasic region is anticipated to bind to anionic phospholipids, including PS, via non-specific electrostatic interactions (4, 5), which we confirmed in protein-lipid overlay experiments (Fig. S7B). Substitution of seven lysines into neutral glutamines in ROP6 C-terminus (ROP6^7Q^) abolished *in vitro* interactions with all anionic lipids (Fig. S7B). In planta, diminishing the net positive charges of ROP6 C-terminus (ROP6^7Q^) or the net negative charge of the PM gradually induced ROP6 mislocalization into intracellular compartments (Fig. S7C-D). To test the functionality of ROP6^7Q^ at the PM, we selected transgenic lines that had strong expression level to compensate for their intracellular localization and therefore have comparable levels of ROP6^7Q^ and ROP6^WT^ at the PM (Fig. S8A and D-F). ROP6^7Q^ mutants were not functional *in planta* (Fig. 6A-B, Fig. S8B-C), even though the 7Q mutations had no impact on ROP6 intrinsic GTPase activity *in vitro* and ROP6-GTP conformation *in vivo* (Fig. S9). We next analyzed the dynamics of mEos-ROP6^7Q^ at the PM of wild-type roots by sptPALM experiments and found that it had the same proportion of immobile molecules than mEos-ROP6^WT^ in *pss1-3*, and that in both cases they were insensitive to auxin (Fig. 6C, Fig. S5E-H). Therefore, impairing PS/ROP6 interaction by either removing PS from the membranes (*pss1* mutant), or by mutating the anionic lipid-binding site on ROP6 (ROP6^7Q^) similarly impacted ROP6 signaling and its auxin-induced nanoclustering.

**Figure 6.**
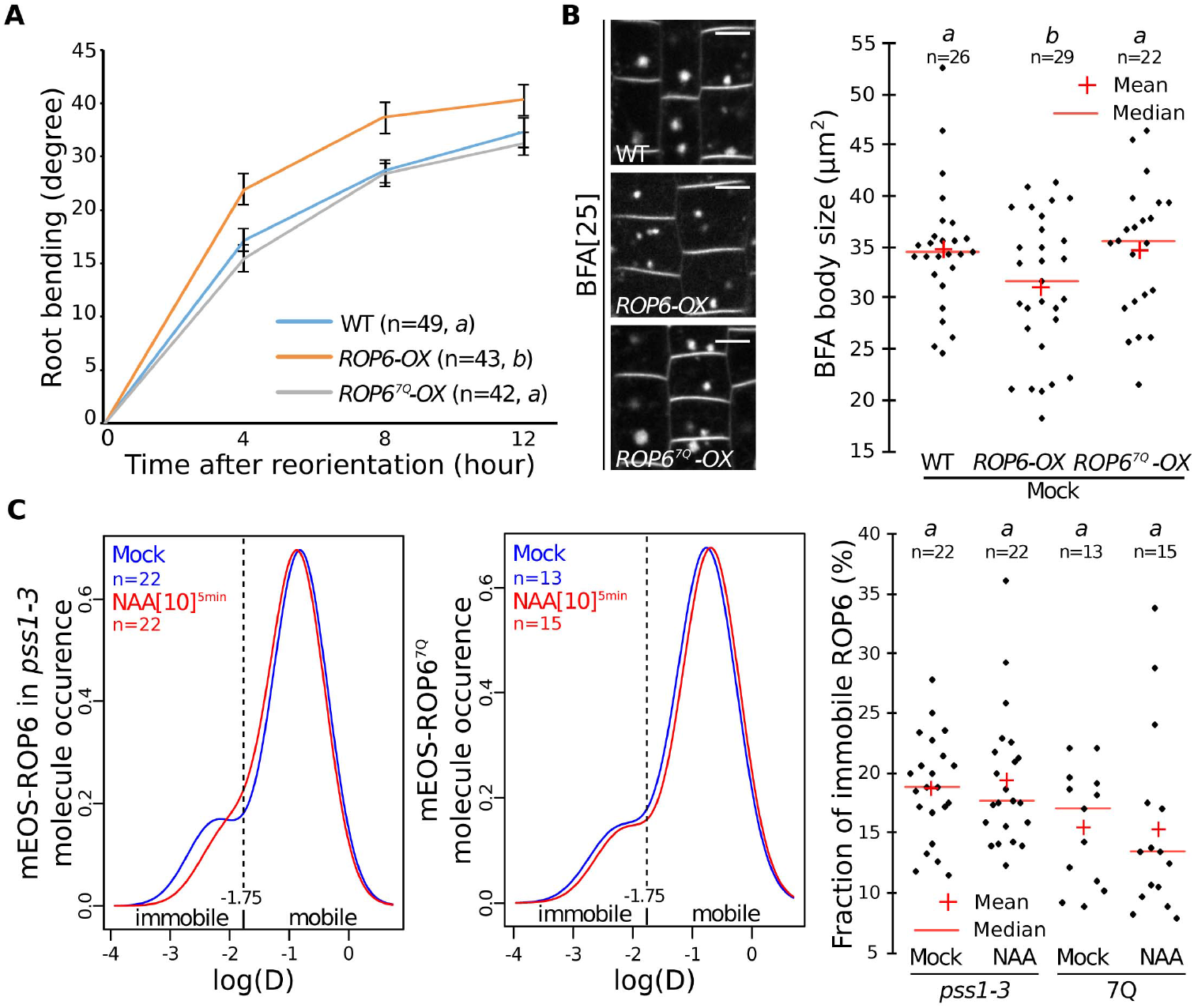
ROP6 nanoclustering in response to auxin requires direct interaction with anionic phospholipids. **A-B**, Quantification of (A) root bending after gravistimulation and (B) the size of FM4-64-stained BFA bodies (data in WT are the same as in Figure 2C). **C**, Distribution of mEos-ROP6 molecules in *pss1-3* (left) and mEos-ROP6^7Q^ molecules in WT (right) roots according to their apparent diffusion coefficient obtained by analysing sptPALM trajectories (NAA: 10μM, 5 min), and related quantification. n represents the number of roots (A, B) or acquisitions (independent cells) (C) analysed, letter indicates statistical difference (see methods for details on statistical tests).

## Conclusions

Here, we showed that in root tip epidermal cells: i) ROP6 is immobilized in PM-nanodomains upon activation by auxin, ii) PS is necessary for both ROP6 stabilization into PM-nanodomains and signaling, iii) ROP6 directly interacts with anionic lipids, including PS, and iv) PS itself is present and immobile in these PM-nanodomains, suggesting that stabilized ROP6 in PS-containing nanoclusters constitutes the functional signaling unit of this GTPase. Our imaging pipeline revealed that ROP6 nano-organization is rapidly remodeled by auxin and as such will provide a quantitative *in vivo* read-out to re-evaluate how auxin may be perceived upstream of ROP6 activation. Given that plants have 11 ROPs, which can respond to a wide range of signals (16), it will be intriguing to address whether nanoclustering is specific to auxin response or common to other signals and to various ROPs, and to what extent it may contribute to signal integration by plant Rho GTPases. All ROP proteins have polybasic clusters at their C-terminus (Fig. S10A), and PS could therefore potentially regulate additional member of this family. Interestingly, in addition to root gravitropic defects, *pss1* had many developmental phenotypes that may be linked to altered ROP function (e.g. pavement cell and root hair morphology, planar polarity defects, see Fig. S10B-F) but that are not found in *rop6* single mutant and could therefore involve additional ROP proteins. Furthermore, nanoclustering seems to be a shared feature of several yeast and animal small GTPases, including K-Ras, Rac1 and Cdc42 (6-8), and both K-Ras and Cdc42 require PS for nanoclustering (3, 6, 8, 11, 13). Here, we found that *in vivo* variations of the PS concentration at the PM act like a rheostat to adjust the sensitivity of ROP6-nanoclustering and hence auxin signaling in a cell-context dependent manner. Our results open the possibility that variations of the PS concentration at the PM in animal systems could also control the signaling capacity of these small GTPases during either normal or pathological development.

## Acknowledgments

We thank M. Dreux, E. Bayer, O. Hamant, S. Mongrand, Y. Boutté, J. Gronnier, J. Reed, T. Vernoux and the SiCE group for discussions and comments, T. Stanislas for help with root hair phenotyping, S. Bednarek, S. Yalovsky, B. Scheres and the NASC collection for providing transgenic *Arabidopsis* lines, A. Lacroix, J. Berger and P. Bolland for plant care, J.C. Mulatier for help in preparing lipids. We acknowledge the contribution of SFR Biosciences (UMS3444/CNRS, US8/Inserm, ENS de Lyon, UCBL) facilities: C. Lionet, E. Chattre, and C. Chamot at the LBI-PLATIM-MICROSCOPY for assistance with imaging and V. Guegen-Chaignon at the Protein Science Facility for assistance with protein purification. We thank the PHIV and MRI platform for access to microscopes.

## Author contributions

M.P.P. generated all transgenic material, and was responsible for all experiments. V.B., M.P.P. and A.M. conceived, performed and analyzed super-resolution imaging. V.B. performed and analyzed TIRFM and FRAP imaging. L.M-P and P.M. performed lipid measurements. A.M. imaged Raichu-ROP6 sensors. J.B. produced recombinant ROP6 and performed GTPase assays. M.P.P and L.A. performed lipid overlay experiments. M.M.M-B., and C.M. assisted with phenotyping and cloning, A.C. performed qRT-PCR analyses, J-B.F. and M.N. designed the sptPALM analyses pipeline, M.P.P., V.B. and Y.J. conceived the study, designed experiments and wrote the manuscript and all the authors discussed the results and commented on the manuscript. Correspondence and requests for materials should be addressed to Y.J.

---

**Figure S1.**
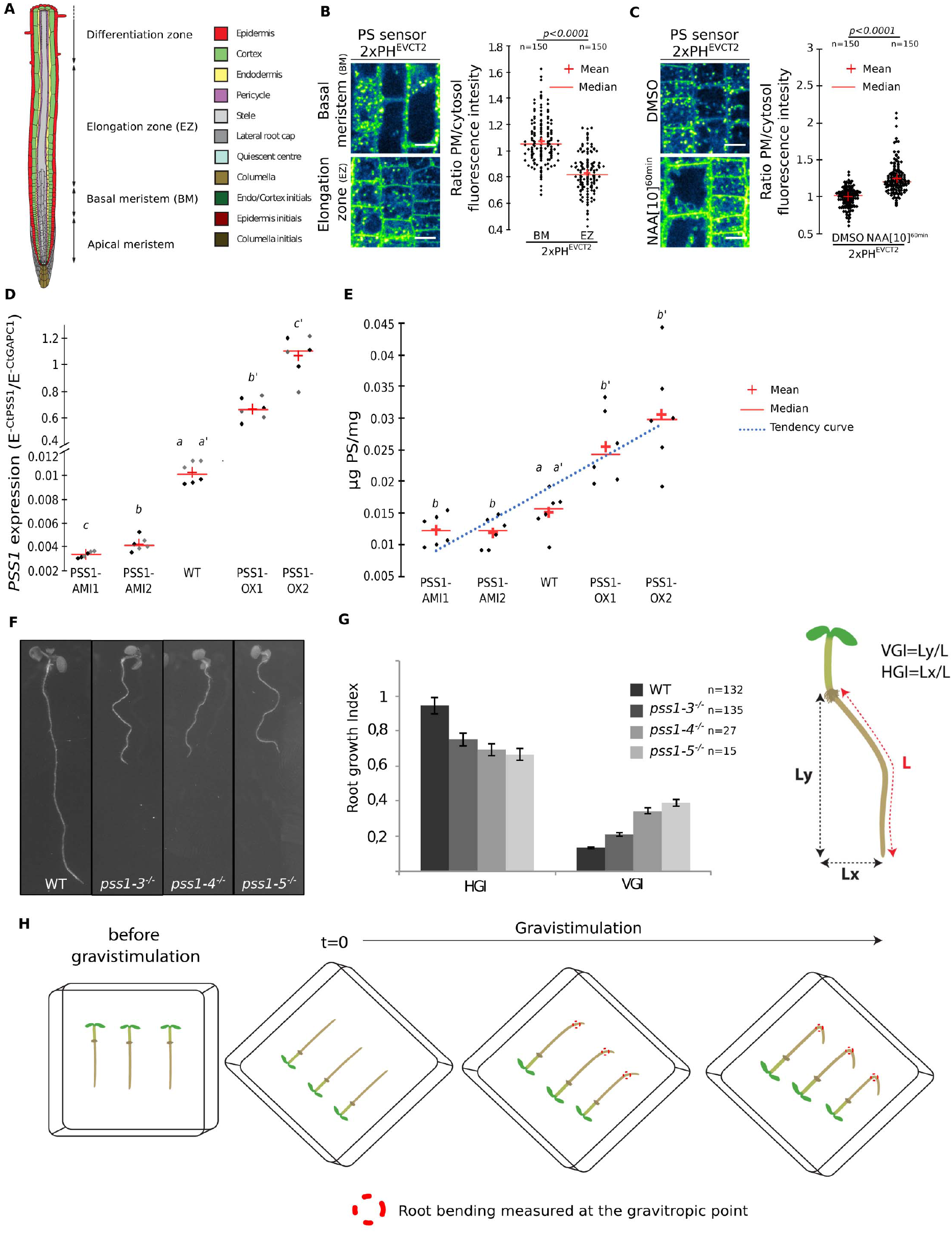
Localization of PS sensor along the root and characterization of lines with graded PS concentration. **A**. Schematic representation of Arabidopsis root with the position of the respective zone of the root. Root picture by Benjamin Peret: https://figshare.com/articles/Primary and lateral root ai/5143987. **B-C**, Confocal images of mCIT-C2^LACT^ root epidermis in basal meristem (BM) and elongation elongation (EZ) in the absence/presence of NAA (10μM, 60min) and related quantification. **D**, qRT-PCR analysis of *PSS1* expression in *WT, PSS1-AMI1, PSS1-AMI2, PSS1-OX1* and *PSS1-OX2* (AMI1/2 are two independent transgenic lines each expressing a different artificial microRNA constructs targeting *PSS1*; *PSS1-OX1/2* are two independent transgenic lines each overexpression *PSS1*). Grey and black dots represent results obtained with different primers used to amplify the *PSS1* cDNA (see Methods). **E**, Quantification of the PS content by HP-TLC in *WT, PSS1-AMI1, PSS1-AMI2, PSS1-OX1* and *PSS1-OX2*. Note that *pss1-3* and *pss1-4* were previously shown to produce no PS (Platre et al., 2018 Dev Cell). **F**, Picture showing the wavy root phenotype of *pss1-3^-/-^*, *pss1-4^-/-^* and *pss1-5^-/-^*, compared to the WT at 12 days after germination (DAG). **G**, Quantification of the horizontal growth index (HGI) and vertical growth index (VGI) in WT, *pss1-3^-/-^*, *pss1-4^-/-^* and *pss1-5^-/-^* at 12 DAG. HGI and VGI were calculated as shown on the cartoon on the right (see Grabov et al., 2005 and methods for details). **H**, Schematic representation of the phenotypic pipeline used to analyse the gravitropic response. A time zero, plants were turned by 135° and root bending at the gravitropic point was analysed using the RootTrace software (see methods for details). n represents the number of cells (B-C) or roots (G) analysed. Scale bars 10μm, letters indicate statistical differences (see methods for details on statistical tests)

**Figure S2.**
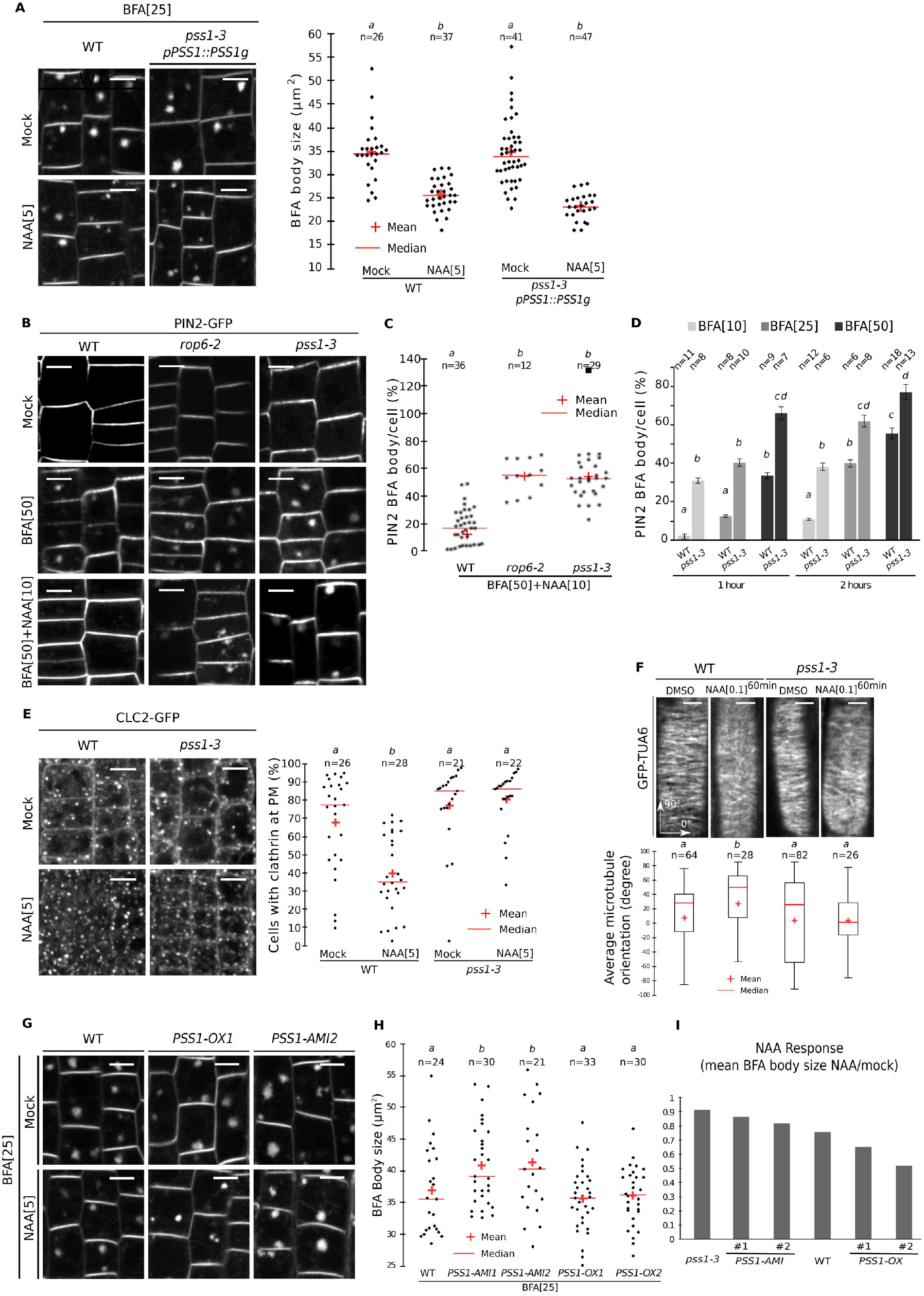
Variations in PS impacts ROP6-mediated auxin responses. **A**, Confocal images of epidermal root cells stained by FM4-64 and treated with BFA (25μM) or BFA/NAA (5μM), and related quantification of the size of FM4-64-stained BFA bodies in WT and complemented *pss1-3/PSS1∷PSS1g* line (BFA: 25μM, 60 min; NAA: 5μM, 30min pre-treatment and 60 min co-treatment with BFA) (image and data in WT are the same as in Fig. 2C). **B**, Confocal images of 10-12-day-old seedlings expressing PIN2prom∷PIN2-GFP in WT, *rop6-2^-/-^, pss1-3^-/-^* (upper panel) and treated with either BFA (50μM, 60 min) or BFA and NAA (10μM, 30 min pre-treatment + 60 min concomitant treatment). **C**, Quantification of the number of PIN2-GFP-positive BFA body number per cell in WT, *rop6-2^-/-^* and *pss1-3^-/-^* concomitantly treated with BFA (50μM, 60 min) and NAA (10μM, 30 min pre-treatment + 60 min concomitant treatment). **D**, Quantification of the number of PIN2-GFP-positive BFA body number per cell in WT and *pss1-3^-/-^* treated with increasing concentration of BFA for one or two hours. **E**, Confocal picture of root epidermal cells expressing CLC2-GFP in the WT and *pss1-3* mutant in the presence or absence of NAA (5μM, 30min) and related quantification of the percentage of cell with visible CLC2-GFP labelling at PM (not considering CLC2-GFP in intracellular compartments). Note that in WT non-treated cells, CLC2-GFP localize at both the PM and intracellular compartments but that auxin selectively depletes the PM pool of CLC2-GFP. **F**, Confocal images of 12-day-old epidermal root cells expressing TUA6-GFP in WT and *pss1-3^-/-^* in presence and absence of NAA (100nM, 60min) and related quantification of the average microtubule orientation. The average orientation was calculated using FibriTool software (see Boudaoud et al., 2014 and methods). **G**, Representative images of confocal micrograph of FM4-64 staining in root epidermis treated with BFA (25μM, 60min, top) or pre-treated with NAA (5μM, 60min) and then co-treated with BFA+NAA (BFA: 25μM, 30min, NAA: 5μM, 60min). Note that the images for the WT are the same as in Fig. 2D. **H**, quantification of the size of FM4-64-stained BFA bodies. **I**, Ratio of the mean size of FM4-64-stained BFA bodies in the NAA and mock conditions (allow to represent the respective NAA response of each line). Scale bars 10μm. n represents the number of roots analysed, letters indicate statistical difference (see methods for details on statistical tests).

**Figure S3.**
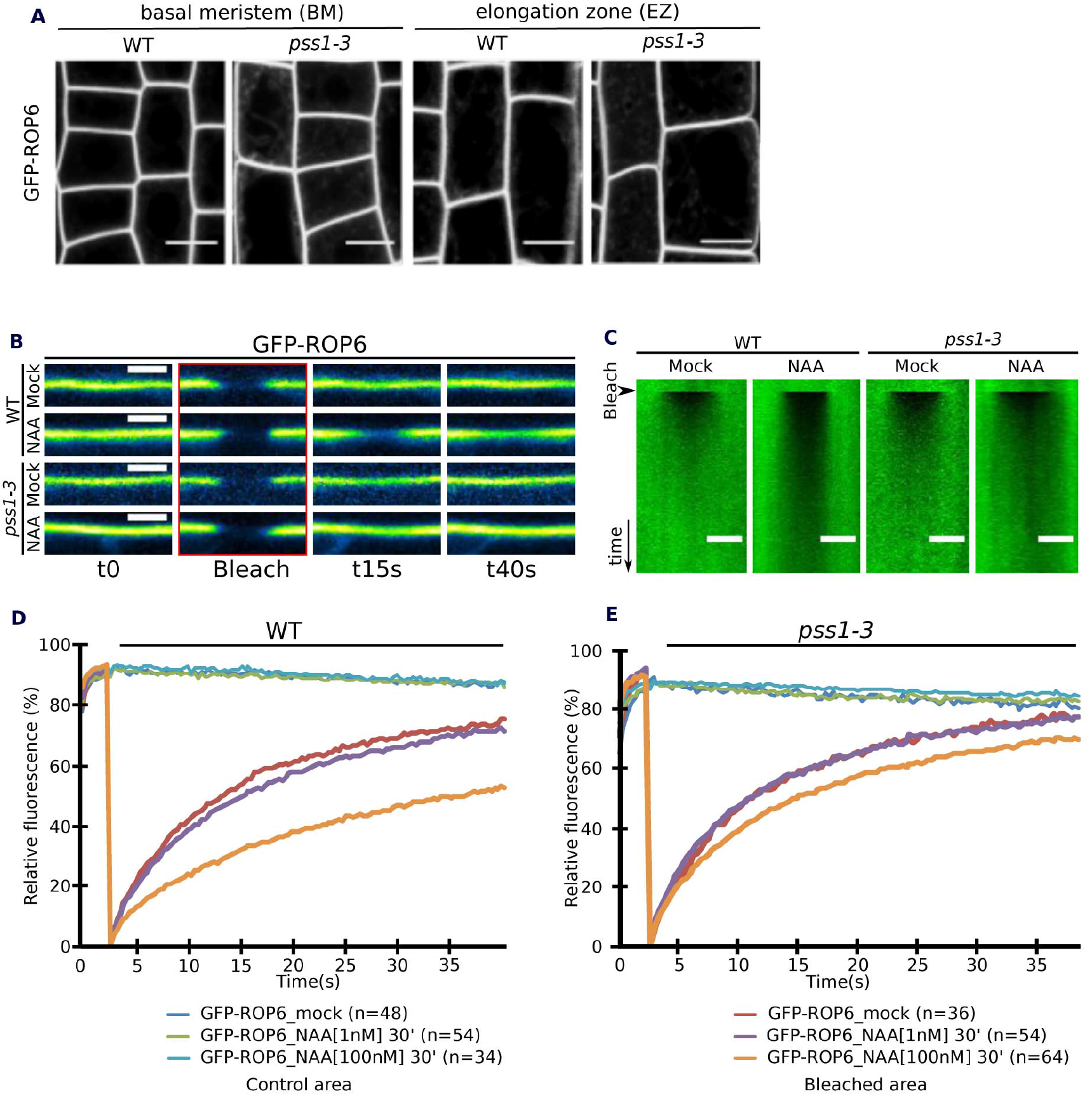
PS is dispensable to efficiently target ROP6 at the PM but is involved in NAA-induced changes in ROP6 lateral diffusion at the PM. **A**, Confocal images of GFP-ROP6 in root epidermis in WT and *pss1-3* mutant in the basal meristem (BM) and elongation zone (EZ). Note that GFP-ROP6 is still efficiently targeted to the PM in *pss1-3*, although we noticed a faint delocalization of GFP-ROP6 in intracellular compartments. Scale bars 10μm. **B**, Confocal images of GFP-ROP6 during FRAP experiment in WT and *pss1-3* in the mock and auxin-treated condition (NAA, 100nM, 30 min), and **C**, related kymograph (time scale 9 seconds). **D-E**, Traces of GFP-ROP6 fluorescence intensity at the PM during FRAP analyses in mock and NAA (30min at 1nm or 100nm) treated conditions in WT and *pss1-3* roots. For each trace in D and E, “n” are similar as in Fig. 4A. Images for the WT are the same as in Fig. 3A and B. Scale bars: 5 μm.

**Figure S4.**
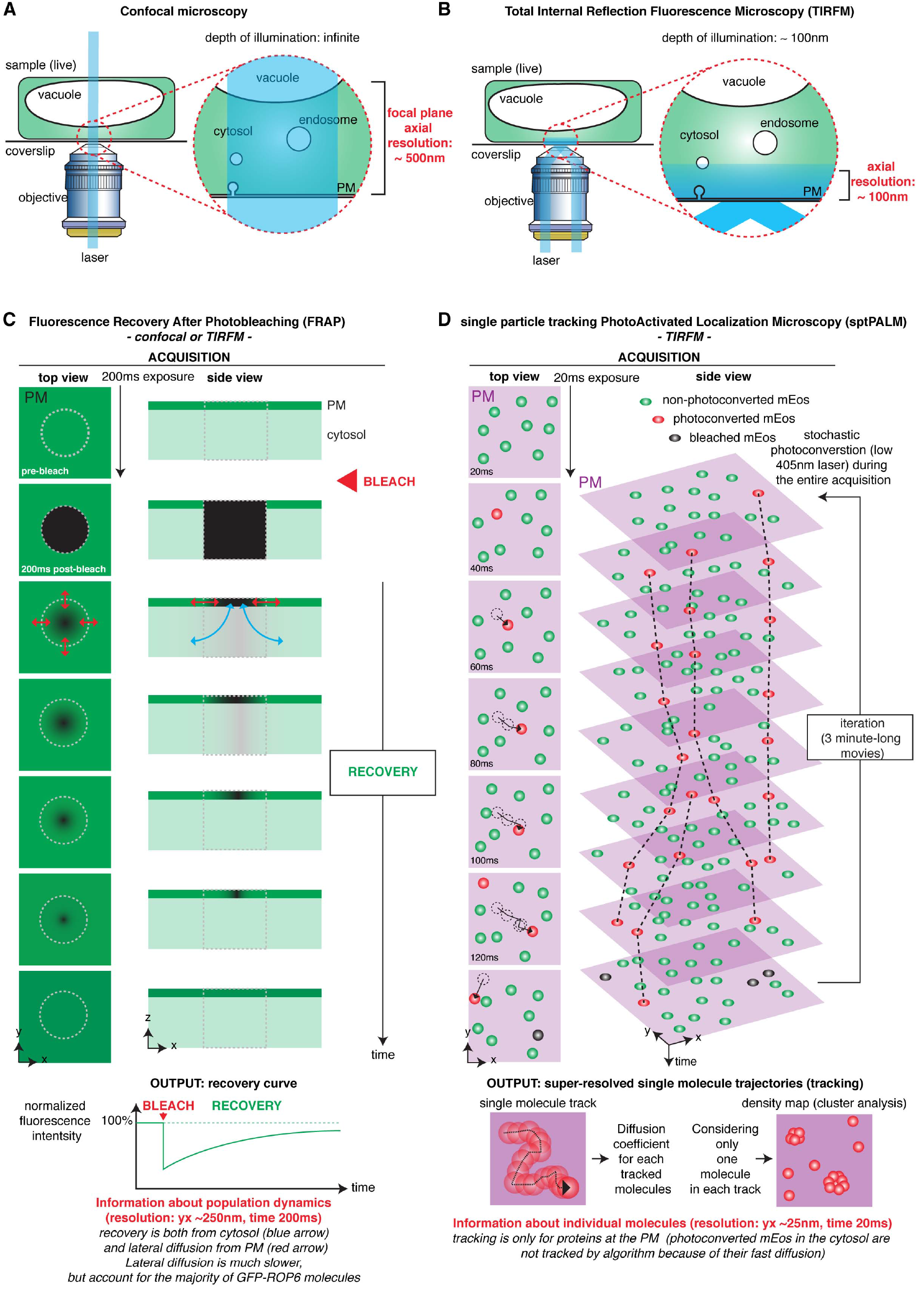
Principle underlying the TIRFM, FRAP and sptPALM analyses. **A**, In confocal microscopy (both laser scanning and spinning-disk confocal microscopy), the illumination beam of the laser is perpendicular to the coverslip and passes through the entire sample, exciting all the fluorophores. The focal plane is determined by the pinhole(s) of the confocal microscope and provides an axial resolution (z position) of about 500nm. **B**, In Total Internal Reflection Fluorescence Microscopy (TIRFM), the illumination beam is oriented at a critical angle when it hits the interface between the coverslip and the sample. In this condition, the beam is reflected, which locally generates an evanescent wave of low energy that penetrates the sample up to about 100nm. As such, only fluorophores that are in close contact with the coverslip are excited, which provides an axial resolution of about the depth of the evanescent wave (~100nm). TIRFM is therefore particularly well suited to study events happening at or in the close vicinity of the PM in contact with the coverslip. Because fluorophores outside of the evanescent wave are not excited, this technique dramatically increases the signal-to-noise ratio during imaging, which allows faster imaging (and/or single molecule imaging). While the xy resolution is still limited (~250nm), the fact that TIRFM largely deplete out-of-focus fluorescence allows to visualise structures, which are in the plane of the PM, and that are not easily identifiable by confocal microscopy. Schematic representation was inspired by Johnson and Vert, 2017. **C**, Fluorescence Recovery After Photobleaching (FRAP) is a technique to study the kinetics of diffusion of fluorescent molecules. It is based on the fact that a focused high intensity illumination will selectively photobleach the fluorophore (i.e. permanently unable to fluoresce) on a determined region of the sample (here, a small portion of the plasma membrane). Through time, the still-fluorescing GFP-ROP6 molecules will diffuse and replace the dark molecules in the bleached region. FRAP provides information about the diffusion dynamics of the population of GFP-ROP6 molecules (i.e. it provides an averaged value of diffusion, but cannot distinguished whether some proteins are diffusing slowly and some others rapidly, or whether their diffusion is uniform). In addition, fluorescence may be recovered from GFP-ROP6 diffusing in the membrane or arriving from the cytosol. Finally, FRAP can be recorded either in confocal microscopy (see Fig. 3A-B and 4A) or TIRFM acquisition (see Fig. 4E). **D**, single particle tracking PhotoActivated Localization Microscopy (sptPALM) is a super-resolution microscopy method, which is a derivative of the PALM technique. PALM uses the stochastic activation of photoactivable (or photoswitchable in the case of mEos) fluorescent proteins to perform single molecule imaging. Photoactivation is performed with low light intensity, so that only a small fraction of the molecules is activated, and therefore photoactivated proteins are well separated from each other during imaging. In order to image single molecules, a microscopy setup with a high signal-to-noise ratio, such as TIRFM, is required. In the framework of our study on ROP6 and PS, TIRFM is an ideal technique since it allowed us to preferentially focus on the pool of ROP6/PS at the PM. Each single molecule is recorded as a diffraction limited spot that can be fitted and thereby positioned with high accuracy (xy resolution of about 25nm). PALM is usually performed on fixed samples and the photoactivated proteins are immediately photobleached during the acquisition (excitation with high laser intensity), so that the position of each molecule is recorded only once. The super-resolved image is built through successive cycle of photoactivation followed by acquisition of the single molecules position and its concomitant photobleaching. The main difference between PALM and sptPALM is that sptPALM is performed on live samples (i.e. with molecules diffusing during the time lapse acquisition) and photoactivated molecules are not immediately photobleached during acquisition (excitation with mild laser intensity). This allows to track the positions of single molecules through time before the photoactivated fluorescent protein is bleached (or detached from the PM). This technique thereby allows to obtain the trajectories of single molecules with a xy resolution of roughly 25nm and time resolution of 20ms. It is then possible to obtain a range of quantitative parameters from these tracks, including a spatial map of the trajectories (as shown on Fig. 3D), or their diffusion coefficient (as shown on Fig. 3E, 5B and 6C). By considering only the position of one molecule per track, we can build “livePALM” images, which shows the average position of each mEos-ROP6 molecule imaged during the acquisition (3-minute-long movies). This provides a density map of mEos-ROP6 and allows us to perform cluster analyses (in our case using Voronoï tessellation, see Fig. 3F-G and 5A). By contrast to FRAP analysis, which provides a global analysis of diffusion at the scale of the bleached fluorescent protein population, sptPALM gives access to single molecules information. Furthermore, sptPALM/TIRFM has also increased resolution in all dimensions (z ~100nm, xy ~25nm, t=20ms vs z ~500nm, xy ~250nm, t=200ms in FRAP/confocal)

**Figure S5.**
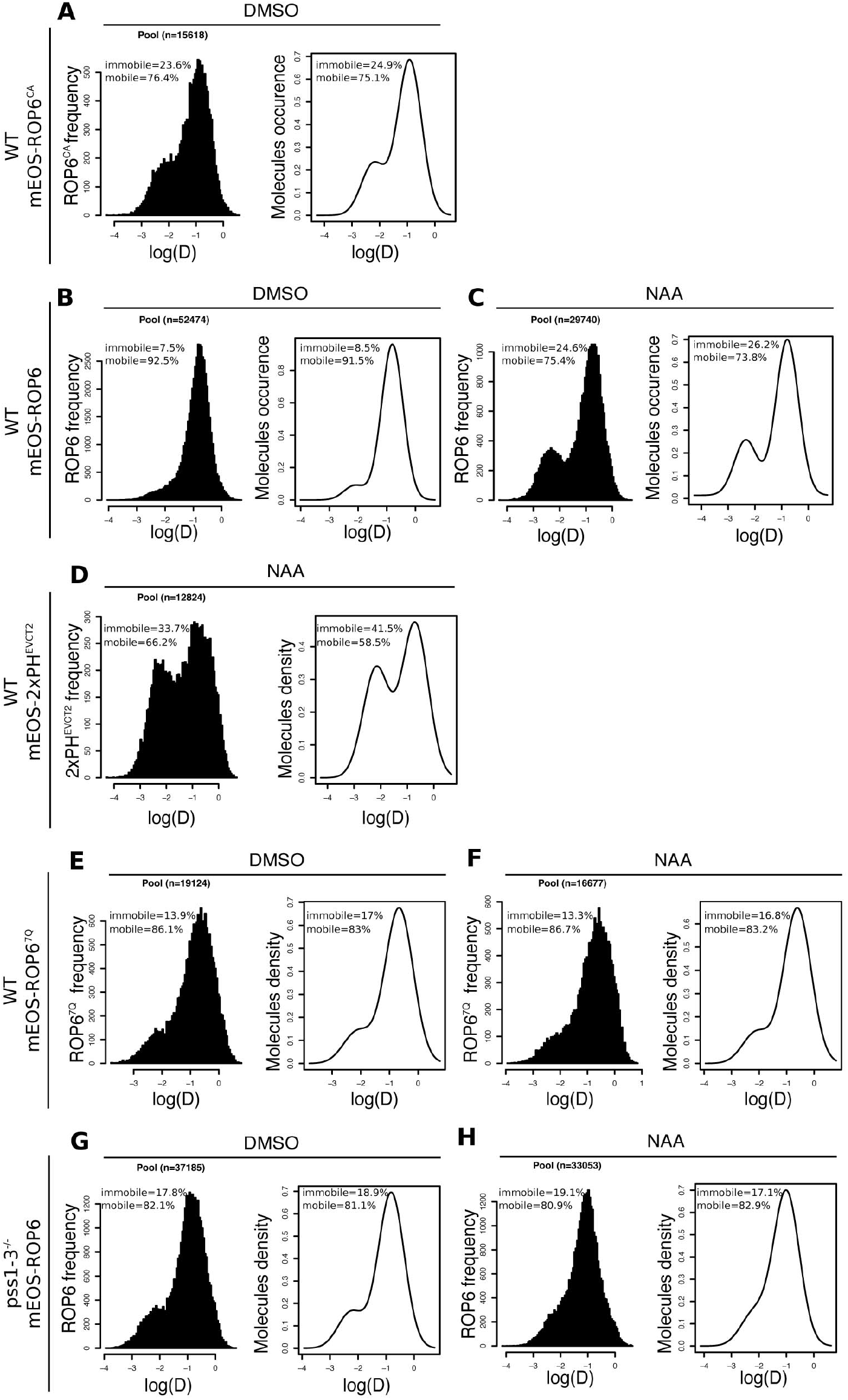
Raw data of sptPALM experiments presented in the paper. **A-H**, Left panel represents the frequency of mEos-ROP6, mEos-ROP6^CA^, mEos-2xPH^EVCT2^, mEos-ROP6^7Q^, mEos-ROP6/*pss1-3* molecules according to their apparent diffusion coefficient (log(D)) obtained by analysing sptPALM trajectories in mock and auxin-treated condition (NAA, 10μM, 5min) in WT or *pss1-3*. Right panel represents the distribution of mEos-ROP6, mEos-ROP6^CA^, mEos-2xPH^EVCT2^, mEos-ROP6^7Q^, mEos-ROP6/*pss1-3* molecules according to their apparent diffusion coefficient (log(D)) obtained by analysing the frequency plot (left panel) using the R mClust package. The percentage of immobile and mobile molecules in each graph represents the percentage of molecules with a trajectory below and above the log(D) of −1.75, respectively. Note that the percentage in left and right panels for each condition are similar (i.e. they vary only from few percent), confirming the existence of two populations with a normal distribution. Note that the right plots in B and C are similar to the plot in Fig. 3E, the right plot in D is similar to the plot in Fig. 5B and the right plots in E-F are similar to the plots in Fig. 6C. “n” corresponds to the number of trajectories analysed

**Figure S6.**
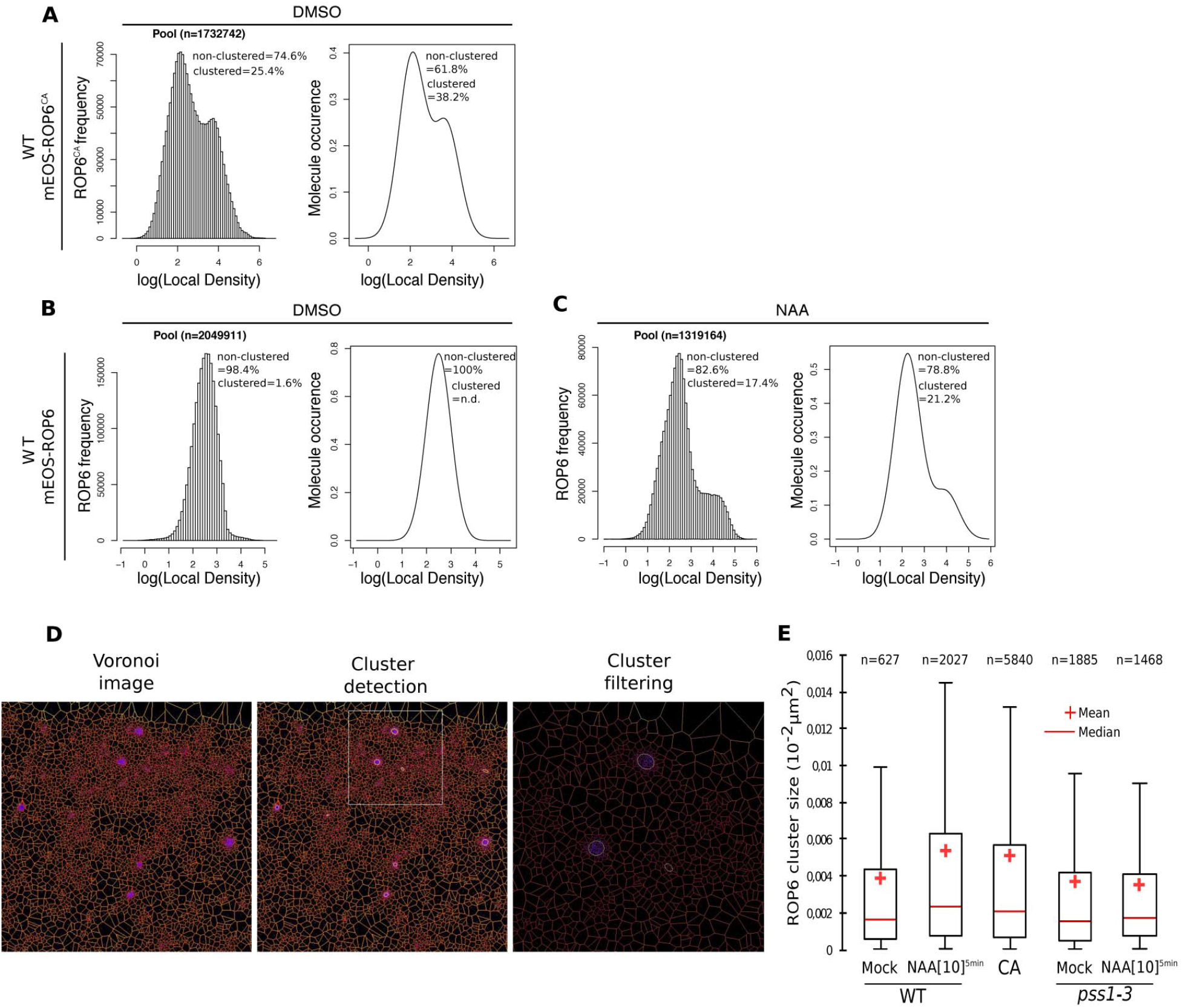
Raw data of live PALM experiments presented in the paper. **A-C**, Left panels represents the frequency of mEos-ROP6 and mEos-ROP6^CA^ molecules according to their local density (log(local density)) obtained by analysing tessellation-based automatic segmentation of super-resolution images in mock and auxin-treated condition (NAA, 10μM, 5min) in WT. Right panel represents the distribution of mEos-ROP6 and mEos-ROP6^CA^ according to their local density (log(local density)) obtained by analysing the frequency plot (left panel) using the R mClust package. The percentage of low and high local density in each graph represents the percentage of molecules below and above the log(local density) of 3.5, respectively. Note that the percentage in left and right panels for each condition are similar (i.e. they vary only from few percent), confirming the existence of two populations with a normal distribution. Note that the right plots in B and C are similar to the plot in Fig. 3G. “n” corresponds to the number of molecules analysed. **D**, Example of Voronoï images obtained after extracting local density from tessellation-based automatic segmentation of super-resolution images (left image), cluster detection and filtering of local density (middle image, zoom in right image) in the Voronoï image and **E**, quantification of the respective size of the cluster in (from left to right), mEos-ROP6 mock-treated, mEos-ROP6 NAA-treated (5min, 10μM), mEosROP6^CA^ mock-treated, mEos-ROP6 mock-treated in *pss1-3* and mEos-ROP6 NAA-treated in *pss1-3* (5min, 10μM). “n” corresponds to the number of nanocluster analysed.

**Figure S7.**
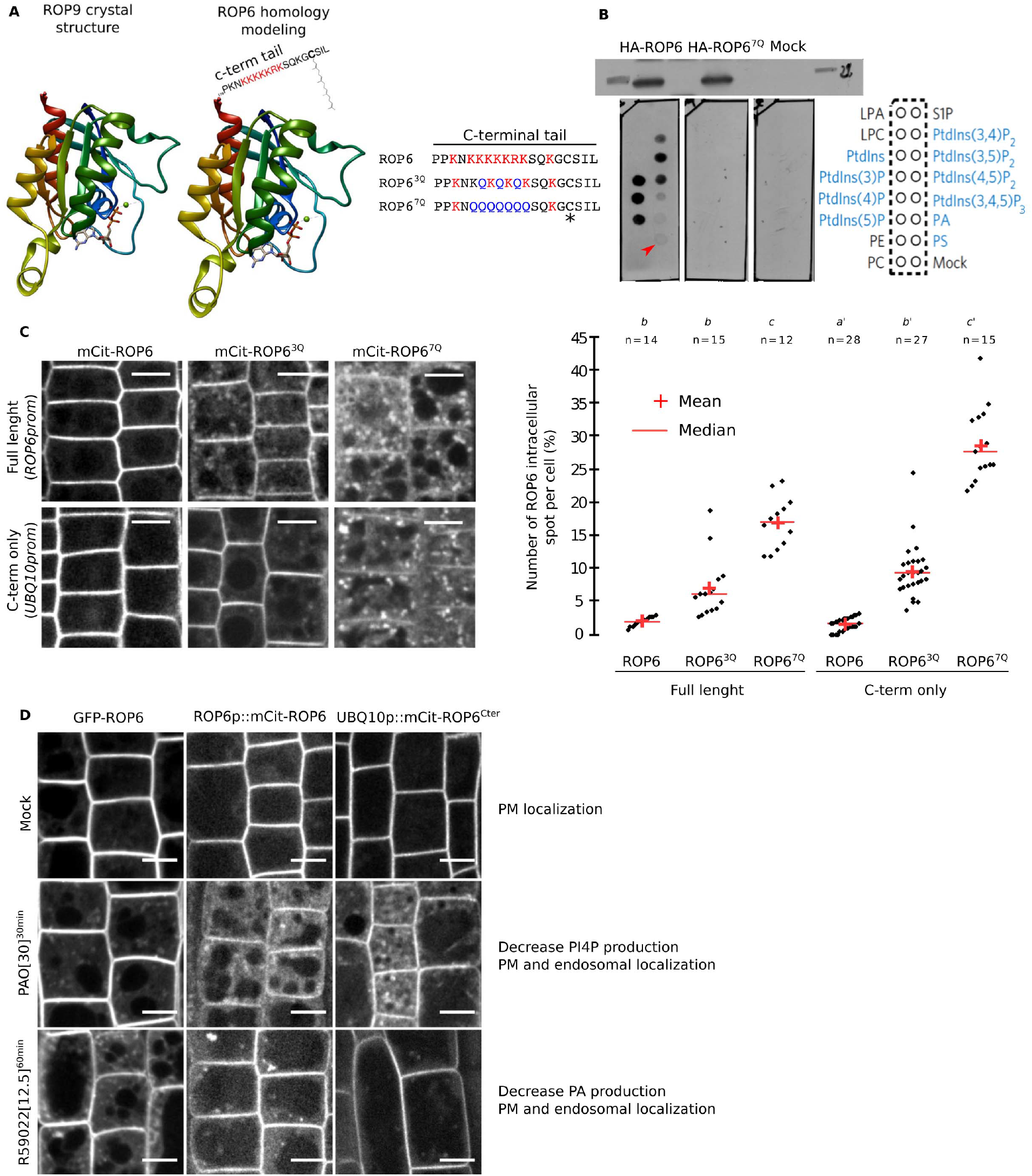
ROP6 polybasic region interacts with anionic phospholipids and is involved in PM targeting. **A**, Structure of ROP9 (PDB: 2J0V) and homology model of ROP6 based on ROP9 structure. The position of ROP6 C-terminal tail (polybasic region + geranylgeranyl) is indicated as well as the sequence of ROP6, ROP6^3Q^, ROP6^7Q^ C-terminal tail. The asterisk indicates the cysteine required for geranylgeranylation. **B**, Western blot (top) showing expression of recombinant HA-ROP6 and HA-ROP6^7Q^ and mock (empty vector) and their corresponding lipid overlay assays (bottom left). Bottom right, scheme showing the position of the different lipid species spotted on the membrane. Anionic lipids are highlighted in blue. The red arrowhead indicates the interaction between HA-ROP6 but not HA-ROP6^7Q^ with PS. **C**, Confocal images of mCIT-tagged *ROP6, ROP6^3Q^, ROP6^7Q^, ROP6^Cter^, ROP6^3Q-Cter^, ROP6^7Q-Cter^* and related quantification of the number of intracellular compartments per cells. n represents the number of roots analysed, letters indicate statistical differences (see methods for details on statistical tests). **D**, Confocal images of *35Sprom∷GFP-ROP6*, *ROP6prom∷mCIT-ROP6* and *UBQ10-mCIT-ROP6* in the mock, PAO (30μM, 30min, PI4-Kinase inhibitor) and R599022 (12.5μM, 60min, DAG-Kinase inhibitor) treated condition. Consistent with ROP6 polybasic region being able to interact with anionic phospholipids via non-specific electrostatic interactions, ROP6 interacted with all anionic phospholipids. In addition, in this *in vitro* assay, ROP6binding to PS and PA was weaker than binding to phosphatidylinositol phosphate (PIPs). This is expected as both PA and PS are less electronegative than PIPs. Nonetheless, this may not reflect the relative importance of these interactions *in vivo*, since they have different relative concentrations. We noticed that the C-terminal tail of ROP6 closely resemble the C-terminal tail of K-Ras that we recently used as a membrane surface charge sensor (MSC) in plants (Simon, Platre et al., 2016; Platre et al., 2018). ROP6, like MSC sensors, possess in its C-terminus a polybasic region (PBR) adjacent to a prenylation site (i.e. geranylgeranylation). Substitution of seven lysine residues into neutral glutamine in ROP6-PBR (ROP6^7Q^) abolished *in vitro* interaction with all anionic lipids, showing that interaction with anionic phospholipids was fully dependent on the positive charges in ROP6 C-terminal tail. In planta, diminishing the net positive charges of ROP6-Cter gradually increased its mislocalization into intracellular compartments. This effect was charge dependent as mutating 3 lysine residues in the polybasic region (ROP6^3Q^) had an intermediate effect on ROP6 PM targeting. Again, the graded delocalization of mutant ROP6 into intracellular compartments depending on their charges closely resembled the behavior of MCS reporters (Simon, Platre et al., 2016; Platre et al., 2018). This suggested that ROP6 PM targeting was dependent on the electrostatic potential of this membrane. Furthermore, we obtained similar results when we expressed only the C-terminal tail of ROP6 (PBR + geranylgeranylation site). Therefore, ROP6 C-terminal tail is necessary and sufficient for ROP6 PM targeting. As shown in Figure S3A, PS has only a minor role in ROP6 PM targeting, with most of GFP-ROP6 remaining at the PM in the *pss1* mutant and only a small proportion being delocalized in intracellular compartments. This was in stark contrast with the extensive delocalization of mCIT-ROP6^7Q^ in intracellular compartments. As ROP6^7Q^ impaired interaction with all anionic lipids, these results suggested that interaction with additional anionic lipids are required for efficient PM targeting of ROP6. We previously demonstrated that three anionic lipids are involved in the plant PM high electrostatic signature, with PI4P playing a major role, followed by PA and then PS (Simon, Platre et al., 2016; Platre et al., 2018). To test whether PI4P and/or PA could be involved in ROP6 PM targeting, we inhibited their synthesis using inhibitors of PI4Kinases (i.e. PAO) and DAG Kinases (i.e. R59022). Again, we found that ROP6 localization (either full length or C-terminal tail only) behaved like MSC reporters, with inhibition of PI4P synthesis having the strongest effect on ROP6 PM targeting followed by inhibition of PA synthesis. Together, these results suggest that ROP6 is targeted to the PM via direct electrostatic interactions between the polybasic region present in ROP6 C-terminal tail and the inner electrostatic potential of the PM. Together, these results also confirm that ROP6/PS interaction main functions are not PM targeting but rather suggest a specific role for this interaction in regulating ROP6 lateral segregation within the PM.

**Figure S8.**
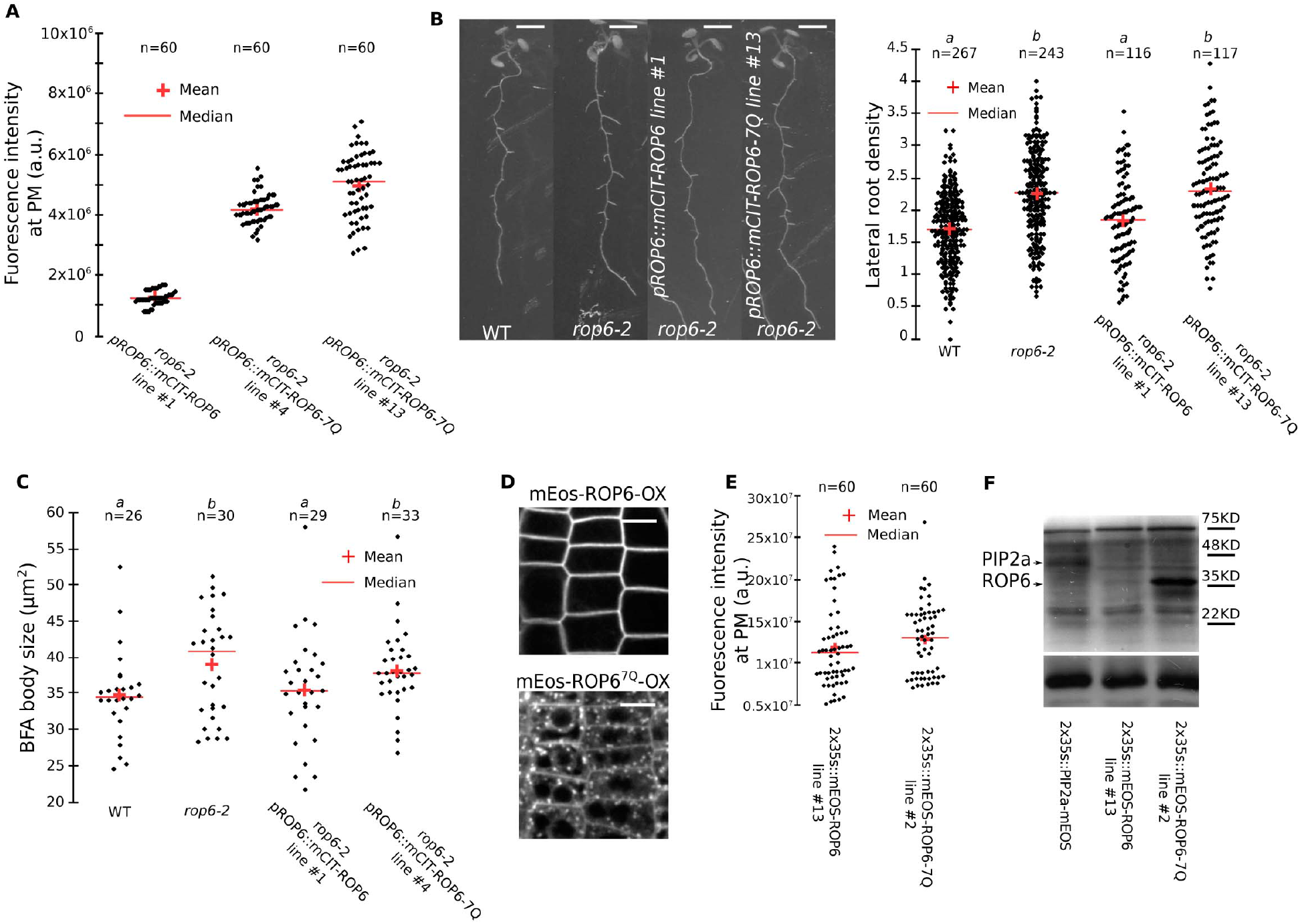
The ROP6 polybasic region is required to complement *rop6-2* loss-of-function allele and to induce *ROP6-OX* gain-of-function phenotype. Because ROP6^7Q^ is delocalized in intracellular compartments (a property that is largely regulated by PI4P and PA, rather than PS, see Fig. S7), we selected transgenic lines with high expression level so that they had similar (or even higher) accumulation at the PM. As expected, we had to select transgenic lines that expressed overall higher amount of proteins than mEos-ROP6 (as seen using anti-mEos western blot analyses) to have similar amount or mEos-ROP6^7Q^ at the PM. This strategy allowed us to perform complementation analyses and to analyse whether PM-localized ROP6^7Q^ is functional or not.**A**, Quantification of the mCITRINE signal at the PM in 7 day-old root meristem expressing *ROP6prom∷mCITRINE-ROP6* and *ROP6prom∷mCITRINE-ROP6^7Q^* in *rop6-2^-/-^* background (n=60 plasma membrane). **B**, Images of 12 day-old seedlings expressing *ROP6prom∷mCITRINE-ROP6* and *ROP6prom∷mCITRINE-ROP6^7Q^* in *rop6-2^-/-^* background showing lateral root formation, and related quantification of the lateral root density. Lateral root formation was used as a sensitive phenotypic read-out for the complementation of *rop6-2* (Lin et al., 2012). **C**, Quantification of BFA body size in *ROP6prom∷mCITRINE-ROP6* and *ROP6prom∷mCITRINE-ROP6^7Q^* in *rop6-2^-/-^* background. **D**, Confocal images of 7 day-old root cells overexpressing *2×35S∷mEos-ROP6 and 2×35S∷mEos-ROP6^7Q^*. **E**, Quantification of the mEos signal at the plasma membrane (integrated intensity) in 7 day-old root meristem overexpressing *2×35S∷mEos-ROP6* and *2×35S∷mEos-ROP6^7Q^*. **F**, Western blot showing mEos-tagged protein accumulation in the following transgenic lines (from left to right): *PIP2a∷PIP2a-mEos, 2×35S∷mEos-ROP6* (line 13) and *2×35S∷mEos-ROP6^7Q^* (line 2). In the top panel, the blot was probed with an anti-mEos antibody and in the bottom panel with anti-Histone H3 antibody as loading control. n represents the number of roots analysed, letters indicate statistical differences (see methods for details on statistical tests).

**Figure S9.**
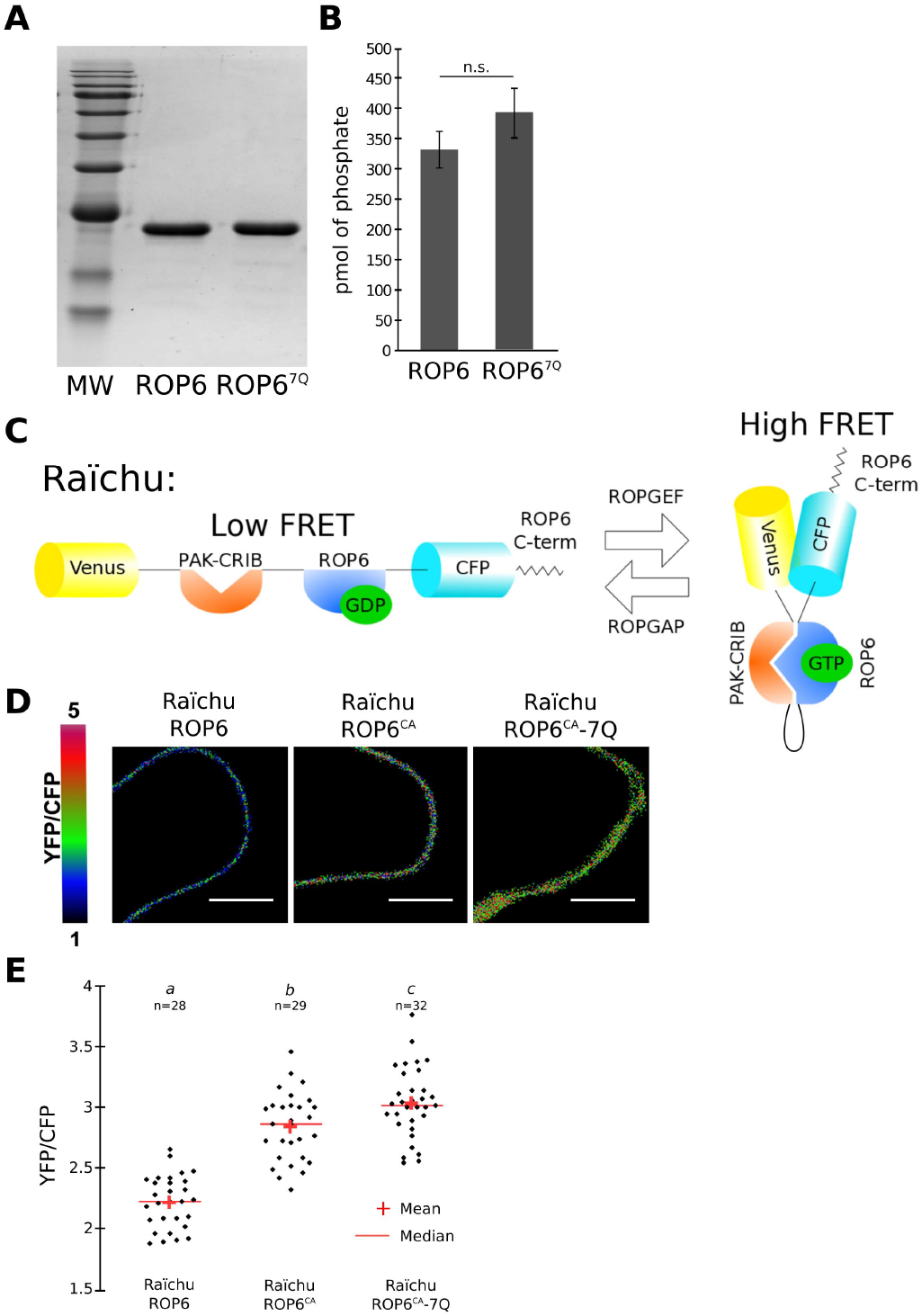
ROP6 polybasic region is not required for its GTPase activity or for interaction with effectors. **A**, SDS-PAGE of purified recombinant 6His-ROP6 (full length) and 6His-ROP6^7Q^. **B**, Measurement of ROP6 and ROP6^7Q^ intrinsic GTPase activity. **C**, Schematic representation of the construct architecture and principle of the FRET-based sensor of ROP6 activation (RaichuROP6). **D**, Ratiometric images of Raichu-ROP6, Raichu-ROP6^CA^, and Raichu-ROP6^CA-7Q^ transiently expressed in *Nicotiana tabacum* (colour scale shown on the left) and **e**, related quantification. n represents the number of cell analysed, letters indicate statistical differences (see methods for details on statistical tests). After having established that PM-localized ROP6^7Q^ are not functional, we verified that mutating the polybasic region did not impacted ROP6 GTPase activity and/or its ability to bind downstream effectors when in a GTP-bound conformation. For this purpose, we purified full length ROP6 (including the entire C-terminal tail with or without the 7Q mutation) and tested its intrinsic GTPase activity (i.e. ability to hydrolyse GTP in solution). This assay showed that the 7Q mutation in ROP6 C-terminal tail did not impacted the enzymatic activity of the GTPase domain. We also developed a ratiometric FRET-based sensor of ROP6 activity based on the design of the Rho GTPase biosensors for Rac/Cdc42 named Raichu-Rac1 and Raichu-Cdc42 (Itoh et al., 2002; Yoshizaki et al., 2003). In Raichu-Rac1/Cdc42 sensors, the CRIB domain of human PAK1 is cloned in tandem with Rac1 or Cdc42 (interspaced by an appropriate linker) and flanked by two fluorescent protein FRET pairs (Venus and ECFP). The CRIB domain interacts specifically with activated Rac1/Cdc42 (i.e. GTP-bound Rac1 or Cdc42), which induces a conformational change in the sensor and enhances the FRET efficiency between the two fluorescent proteins. We used a similar design for the Raichu-ROP6 sensor. Because the CRIB domain of human PAK1 is known to also interact with plants GTP-bound ROPs (Tao et al., 2002; Akamatsi et al., 2013), we used this domain as a generic probe for ROP6-GTP conformation. Transient expression in *Nicotiana tabacum* leaves confirmed that Raichu-ROP6^CA^ had a higher FRET ratio than Raichu-ROP6. We next analysed Raichu-ROP6^CA-7Q^ and found that it has a similar (or even slightly higher) FRET ratio than Raichu-ROP6^CA^. This assay showed that even when the polybasic region in ROP6 C-terminal tail is mutated, ROP6 is still able to interact with a downstream generic effector such as the CRIB domain of PAK. Although it is impossible to test the effect of the 7Q mutations on all the possible effectors of ROP6, this analysis suggests that the 7Q mutations do not impact the overall conformation of ROP6-GTP. Together with the fact that ROP6^7Q^ is still an active GTPase *in vitro* and that the polybasic region is outside of the GTPase domain of ROP6, our analyses strongly suggest that the 7Q mutations do not impact the folding/conformation of ROP6 and rather specifically affects the interaction between ROP6 and anionic phospholipids.

**Figure S10.**
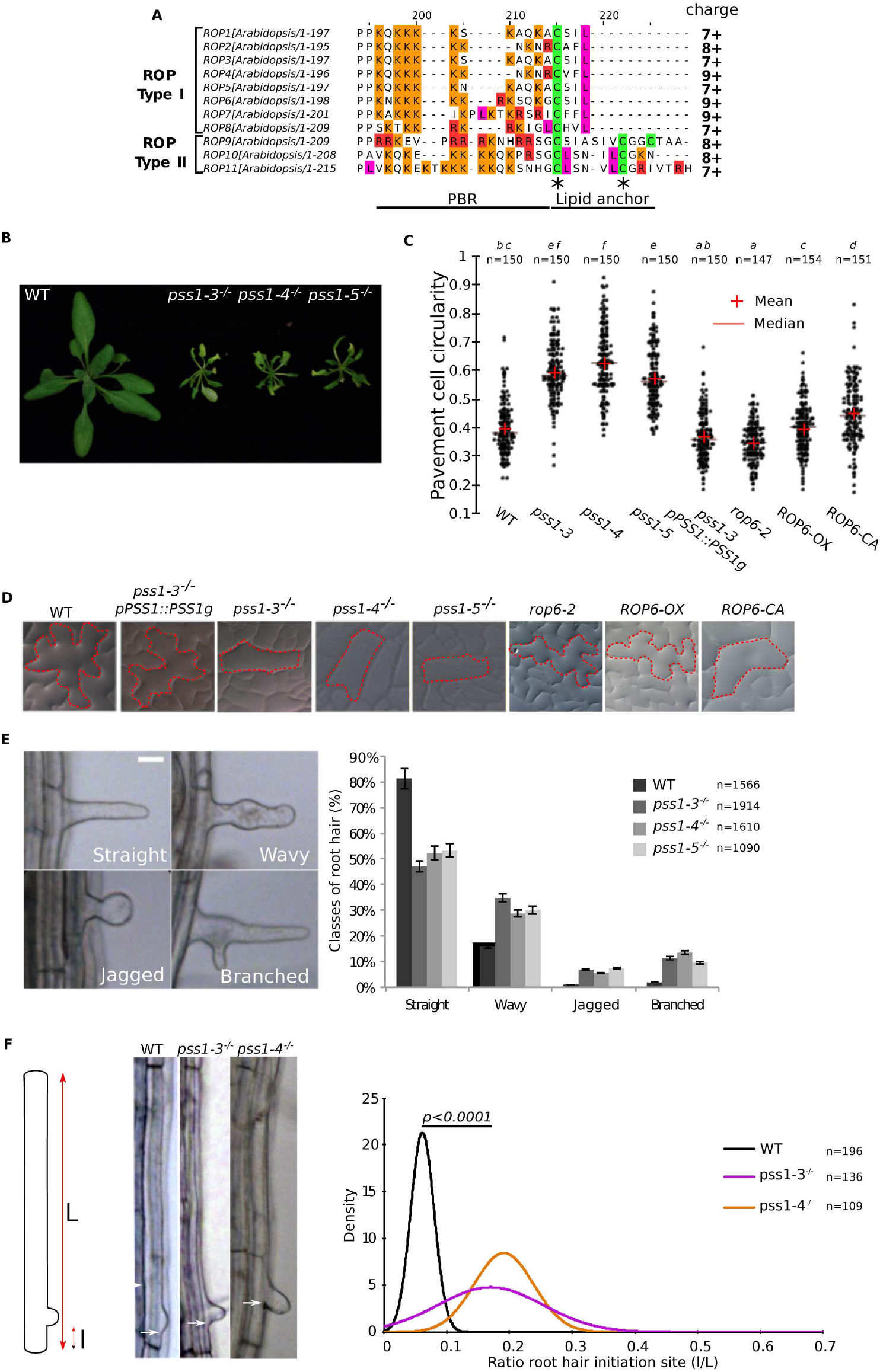
PS-less mutants share ROPs-associated phenotypes. **A**, Sequence alignment of the C-terminal tail of the 11 Arabidopsis ROPs. Note that they all have a polybasic region with net positive charges ranging from +7 to +9. **B**, Pictures of WT, *pss1-3^-/-^*, *pss1-4^-/-^* and *pss1-5^-/-^* rosettes at 21 days after germination (DAG). **C**, Quantification of the pavement cells circularity of WT, *pss1-3^-/-^*, *pss1-4^-/-^*, *pss1-5^-/-^*, *pss1-3^-/-^*x*pPSS1∷PSS1g*, *rop6-2^-/-^*, *ROP6-OX (35S∷GFP-ROP6*) and *ROP6-CA (35S∷GFP-ROP6^CA^*). **D**, Picture showing pavement cells shape of WT, *pss1-3^-/-^*, *pss1-4^-/-^*, *pss1-5^-/-^*, *pss1-3^-/-^*x*pPSS1∷PSS1g*, *rop6-2^-/-^*, ROP6-OX and ROP6-CA. **E**, Picture of representative root hair shape phenotypes observed in *pss1* mutants (classified as straight, wavy, bulged and branched), and related quantification. **F**, Representation of root hair initiation site ratio calculation and picture of WT, *pss1-3^-/-^* and *pss1-4^-/-^* root hair initiation site, and related quantification. Arrows indicate the root hair initiation site. n represents the number of cells (C), and root hairs (E and F) analysed, letters indicate statistical differences (see methods for details on statistical tests).

**Supplementary Video 1:** Example of video showing mEos-ROP6 single molecule imaging used for sptPALM and livePALM analysis.

**Supplementary Video 2:** Video showing GFP-ROP6 localization (10μM NAA, 20 min) in TIRFM before and after bleaching a region of interest containing GFP-ROP6 nanoclusters. Note that GFP-ROP6 nanoclusters are immobile and that they do not recover fluorescence after photobleaching (by contrast to surrounding “non-clustered” GFP-ROP6 signal at the PM, for which recovery of fluorescence is fast)

**Supplementary Video 3:** Video showing GFP-ROP6 localization (10μM NAA, 20 min) in *pss1-3* mutant in TIRFM before and after bleaching a region of interest containing GFP-ROP6 nanoclusters. Note that, like in the WT, GFP-ROP6 nanoclusters are immobile. However, by contrast to the WT, GFP-ROP6 nanoclusters rapidly recover fluorescence after photobleaching.

## Supplementary materials

### Materials and methods

#### Growth condition and plant materials

*Arabidopsis thaliana* Col-0 accession was used as wild type (WT) reference background throughout this study. Plants were grown in soil under long-day conditions at 21°C and 70% humidity and *in vitro* on Murashige and Skoog (MS) Basal Medium supplemented with 0.8% plant agar (pH 5.7) in continuous light conditions at 21°C. Every plant used for experiments are homozygous lines or F2 crosses. *Nicotiana tabacum* cv. Petit Havana plants were grown and used in transient transformation experiments as described (38). The following lines were published before: *pss1-3, pss1-4, pss1-5, pss1-3xPSS1prom∷PSS1genomic; UBQ10prom∷mCHERRY-C2^Lact^ (NASC# N2107778), UBQ10prom∷mCITRINE-2xPH^EVCT2^ (NASC# N2107779*) (28)*;UBQ10prom∷mCITRINE-C2^Lact^ (NASC# N2107347*) (30)*;35Sprom∷GFP-ROP6, 35Sprom∷GFP-ROP6^CA^* (34)*;rop6-2* (22)*;CLC2prom∷CLC2-GFP* (39)*;35S∷GFP-TUA6 (NASC# N6551);PIN2prom∷PIN2-GFP* (40)*;UBQ10prom∷PIP2a-mEos* (15).

#### Microscopy setup

All imaging experiments were performed with the following spinning disk confocal microscope set up, except when indicated otherwise (see bellow): inverted Zeiss microscope (AxioObserver Z1, Carl Zeiss Group, http://www.zeiss.com/) equipped with a spinning disk module (CSU-W1-T3, Yokogawa, www.yokogawa.com) and a ProEM+ 1024B camera (Princeton Instrument, http://www.princetoninstruments.com/) using a 63x PlanApochromat objective (numerical aperture 1.4, oil immersion). GFP was excited with a 488nm laser (150mW) and fluorescence emission was filtered by a 525/50 nm BrightLine^®^ singleband bandpass filter (Semrock, http://www.semrock.com/). YFP/mCITRINE were excited with a 515nm laser (60mW) and fluorescence emission was filtered by a 578/105nm BrightLine^®^ single-band bandpass filter (Semrock, http://www.semrock.com/). mCHERRY and FM4-64 were excited with a 561 nm laser (80 mW) and fluorescence emission was filtered by a 609/54 nm BrightLine^®^ single-band bandpass filter (Semrock, http://www.semrock.com/). 488 or 515 nm lasers were used to excite GFP or YFP/mCITRINE. For quantitative imaging, pictures of epidermal root meristem cells were taken with detector settings optimized for low background and no pixel saturation. Care was taken to use similar confocal settings when comparing fluorescence intensity or for quantification.

Tabaco leaves expressing raichu FRET sensors were observed with an inverted Leica SP8 laser scanning microscope using a 40x Plan-Apochromat objective (numerical aperture 1.1, water immersion). Fluorescent proteins were excited with 458 nm line of a Ne-Ag laser and light were collected simultaneously from 467 to 493 nm for CFP channel and from 538 to 573 for YFP channel. The HyD spectral detector was set to avoid bleed through of CFP fluorescence in the YFP^FRET^ channel. A pinhole of 1 Airy unit was defined. Pixel size and dwell time was respectively 0.143μm and 1.2μs. Those settings were defined to record no signal on nontransformed leaves and kept similar for Raichu constructs tested.

#### FRAP experiments

Fluorescence in a rectangle ROI (50 μm^2^, 15 μm long) was bleached in the plasma membrane optical section by four successive scans at full laser power (150 W) using the iLas2 FRAP module (Roper scientific, http://www.biovis.com/ilas.htm) of our spinning disc microscope (cf description of the system above). Fluorescence recovery was subsequently analysed in the bleached ROIs and in controlled ROIs (rectangle with the same dimension in unbleached area). FRAP was recorded continuously during 90 s with a delay of 0.3 s between frames. Fluorescence intensity data were normalized as previously described (41).

#### TIRF and FRAP experiments in TIRF mode

Total Internal Reflection Fluorescence (TIRF) Microscopy was done using the inverted Zeiss microscope previously described (AxioObserver Z1) equipped with azimuthal-TIRF iLas2 system (Roper Scientific) and a 100x Plan-Apochromat objective (numerical aperture 1.46, oil immersion). Pixel size was 0.13μm. The azimuth calibration was performed using the manufacturer recommendation and the iLas2 module in Metamorph software. The optimum critical angle was determined as giving the best signal-to-noise ratio. Images were acquired with 200ms exposure time, and 300ms between frames in time-lapse experiments. The photobleaching of ROP6 microdomains was achieved on circular ROIs (15μm diameter) by two successive scans at full laser power (150 W) using the iLas2 FRAP module. Pre- and post-acquisitions were recorded in TIRF during 5 minutes with a delay of 0.3 s between frames. In graphs, relative to domain recovery after photobleaching, “n” represents the number of region of interest (ROI) used for quantification.

#### sptPALM

Imaging was performed on a Zeiss Elyra PS1 system with a 100x Apo (numerical aperture 1.46 Oil objective), in TIRF mode. The optimum critical angle was determined as giving the best signal-to-noise ratio. Pixel size was 0.107μm. mEOS was photoconverted using 405nm UV laser power and resulting photoconverted fluorophores were excited using 561nm laser. UV laser power was adjusted to have significant number of tracks without too high density to facilitate further analyses (0.01 to 0.08%). 10,000 images time series were recorded at 50 frames per second (20ms exposure time) on a 256 x 256 pixels region of interest. Single molecule detection and tracks reconstruction were made using MTT algorithm (42) and further computational analyses of tracks were made using CBS sptPALM analyser (Martinière et al., unpublished).

#### FM4-64, BFA, NAA, PAO, R59022 and Lyso-PS treatments

For BFA body size quantification, the plasma membrane and endosomes of 7 to 10-day old transgenic lines were stained by incubating roots with 1 μM FM4-64 (thermofisher scientific, https://www.thermofisher.com) concomitantly with Brefeldin A at 25 μM (BFA, Sigma, www.sigmaaldrich.com, BFA stock solution at 50 mM in DMSO) in liquid MS solution for 60 min. For the auxin analog, Naphthaleneacetic acid (NAA) effect on BFA body size, plants were pretreated with NAA for 30 minutes at 5 μM (Sigma, http://www.sigmaaldrich.com/, NAA stock at 10 mM in DMSO) and then the plasma membrane and endosomes of 5 to 10-day old transgenic lines were stained by incubating roots with 1 μM FM4-64 concomitantly with BFA at 25 μM and NAA at 5 μM in liquid MS solution for 60 min. For NAA effect on microtubule orientation, 10-12-day old seedlings expressing GFP-TUA6 were treated with NAA at 100 nM for 60 minutes. NAA at 5 μM for 30 minutes was applied on 10 to 12-day old seedlings to quantify the percentage of cell showing CLC2 at the plasma membrane. For PIN2 endocytosis, transgenic lines expressing PIN2-GFP (10 to 12 day-old seedlings), were treated with BFA at the indicated time (60 min or 120 min) and concentration (10μM, 25μM or 50μM) in 12-well plates. For NAA effect on endocytosis, PIN2-GFP expressing lines were pretreated with NAA at 10 μM for 30 minutes and then concomitantly treated with NAA at 10 μM and BFA at 50 μM for 60 min in 12-well plates. For NAA effect on PS biosensors *mCITRINE-C2^LACT^* and *mCITRINE-2xPH^EVCT2^*, 5 to 7-day old seedlings were treated with 10 μM NAA for 60 min. 5 to 7 day old seedlings expressing *35Sprom∷GFP-ROP6 (GFP-ROP6*), *ROP6prom∷mCITRINE-ROP6* (mCit-ROP6) and *UBQ10prom∷mCITRINE-ROP6-c-term* (mCit-ROP6^Cter^) were incubated in wells containing 12.5 μM R59022 for 60 min or PAO at 30μM for 30 minutes. Plants observed in TIRFM were treated with NAA at 10μM for 20 minutes. For sptPALM experiment, plants were incubated in DMSO for 20 minutes in 12-well plates and then NAA treatment was performed at 10μM (stock solution 100mM in DMSO) for 5 minutes. For FRAP experiment, NAA was applied at 1nm and 100nm for 30 min. For each treatment, the mock condition corresponds to incubation of plants in well supplemented with a volume of DMSO equivalent to the highest drug concentration used and for the same time as the actual treatment. Roots were imaged within a 5-minute time frame window around the indicated time. Lyso-PS treatment were performed as described (28).

### CLONING

#### Preparation of gateway compatible entry clones (entry vector)

Published gateway compatible entry vectors are listed below.

The *ROP6* promoter (*ROP6prom*) was amplified from Col genomic DNA. Gateway compatible PCR products were introduced into *pDONRP4R-P1* vectors (Thermofisher, www.thermofisher.com, cat# 12537023) by BP recombination using the following primers: *ROP6prom_F* (ttttttgtacaaacttgcctttctctccttcttcaaacttc) and *ROP6prom_R* (gtatagaaaagttgctaacaagctttcagaaaagaggatg) to give *ROP6prom/pDONRP4RP1*.

The ROP6 genomic sequence (*ROP6g – At4g35020*) from the ATG to its 3’UTR was amplified from Col0 genomic DNA. Gateway compatible PCR products were introduced into *pDONRP2R-P3* (Thermofisher, www.thermofisher.com, cat# 12537023) vectors by BP recombination using the following primers: *ROP6-B2R (ggggacagctttcttgtacaaagtggctatgagtgcttcaaggtttatcaagtg*) and *ROP6-B3w3’UTR* (*ggggacaactttgtataataaagttgccttaagacaattggtgtgaatctagg*) to give *ROP6g/pDONRP2RP3*.

Mutation in *ROP6* were obtained by successive site directed mutagenesis using the following partially overlapping forward and reverse primers:

*ROP6-CA-fw* (*gtcggcgacgttgctgttggaaagacttgtc*) and *ROP6-CA-Rev* (*tccaacagcaacgtcgccgacagtgacacacttgataaacc*) using *ROP6g/pDONRP2RP3* as template to give *ROP6g-CA/pDONRP2RP3*.

Mutations in *ROP6g-PBR* were obtained by ligation using the following 5’-phosphorylated primers: *ROP6g-7Q_F* (*gctgctgctggttttttggtggctggagaacgac*) and *ROP6g-7Q_R* (*agcagcaacaatctcagaaaggttgttctatactc*) using *ROP6g/pDONRP2RP3* as template to give *ROP6g^7Q^/pDONRP2RP3.ROP6g-3Q_F* (*ccaaaaaacaagcagaagcagaagcagaaatctcagaaaggttgttc*) and *ROP6g-3Q_R* (*gagatttctgcttctgcttctgcttgttttttggtggctggagaacgacc*) using *ROP6g/pDONRP2RP3* as template to give *ROP6g^3Q^/pDONRP2RP3*. The *ROP6* coding sequence (CDS) was amplified from Col0 cDNA. Gateway compatible PCR products were introduced into *pDONRP221* (Thermofisher, www.thermofisher.com, cat# 12536017) vectors by BP recombination using the following primers: *ROP6-B1* (*ggggacaagtttgtacaaaaaagcaggcttaatgagtgcttcaaggtttatcaagtg*) and *ROP6-B2wSTOP* (*ggggaccactttgtacaagaaagctgggtatcagagtatagaacaacctttctgag*) to give *ROP6cDNA/pDONRP221*.

Mutations in *ROP6cDNA-PBR* were obtained by ligation using the *ROP6g-7Q_F* and *ROP6g-7Q_R* 5’ phosphorylated primers and *ROP6cDNA/pDONRP221* as template to give *ROP6^7Q^cDNA/pDONRP221*.

The *ROP6 C-*terminal tail, wild type and mutated ones were generated using the following 5’phosphorylated primers followed by a ligation:

*ROP6C-term_F* (*aaaatctcagaaaggttgttctatactctaagcaactttattatacaaagttggc*) and *ROP6Cterm_R* (*gctgctgctggttttttggagccactttgtacaagaaagttgaacg*) using *mCITRINE/pDONRP2RP3* as template to give *ROP6-C-term/pDONRP2RP3*.

*ROP6-3Q-C-term_F* (*agaaatctcagaaaggttgttctatactctaagcaactttattatacaaagttggc*) and *ROP6-3QCterm_R* (*gcttctgcttctgcttgttttttggagccactttgtacaagaaagttgaacg*) using *mCITRINE/pDONRP2RP3* as template to give *ROP6^3Q^-C-term/pDONRP2RP3*.

*ROP6-7Q-C-term_F* (*agcagcaacaatctcagaaaggttgttctatactctaagc*) and *ROP6-7QCterm_R* (*gctgctgctggttttttggagccactttgtacaagaaagttgaacg*) using *mCITRINE/pDONRP2RP3* as template to give *ROP6^7Q^-C-term/pDONRP2RP3.mEos/pDONRP221* was obtained by amplifying *mEos* using the following primers: *mEOS_F* (*ggggacaagtttgtacaaaaaagcaggcttaatgagtgcgattaagccagacatgaag*) and *mEOS_R(ggggaccactttgtacaagaaagctgggtattatcgtctggcattgtcaggcaatc*) followed by BP cloning into *pDONRP221*.

The *PSS1* (*At1g15110*) coding sequence was amplified from Col0 cDNA. Gateway compatible PCR products were introduced into *pDONRP221* vectors by BP recombination using the following primers: *PSS1-OX_F* (*ggggacaagtttgtacaaaaaagcaggcttaaccatggaacccaatgggtacaggaaa*) and *PSS1-OX_R(ggggaccactttgtacaagaaagctgggtaacgtctcttttgcgcgaggatcttct*) to give *PSS1cDNA/pDONRP221.PSS1* artificial microRNAs were generated using WMD3-Web MicroRNA designer (Ossowski Stephan, Fitz Joffrey, Schwab Rebecca, Riester Markus and Weigel Detlef, personal communication). The *PSS1-AMI1_B1_B2* and *PSS1-AMI2_B1_B2* were produced by IDT to be introduced into *pDONRP221* vectors by BP recombination.

*PSS1-AMI1_B1_B2: Acaagtttgtacaaaaaagcaggctcaaacacacgctcggacgcatattacacatgttcatacacttaatactcgctgttttgaattgatgttt taggaatatatatgtaga**taataatgatgcgcttaacgt**tcacaggtcgtgatatgattcaattagcttccgactcattcatccaaataccgag tcgccaaaattcaaactagactcgttaaatgaatgaatgatgcggtagacaaattggatcattgattctctttga**acgttaagcgcatcattat ta**tctctcttttgtattccaattttcttgattaatctttcctgcacaaaaacatgcttgatccactaagtgacatatatgctgccttcgtatat atagttctggtaaaattaacattttgggtttatctttatttaaggcatcgccatgacccagctttcttgtacaaagtggt*

*PSS1-AMI2_B1_B2: acaagtttgtacaaaaaagcaggctcaaacacacgctcggacgcatattacacatgttcatacacttaatactcgctgttttgaattgatgttt taggaatatatatgtaga**tttaacgtctcttttgcgcgc**tcacaggtcgtgatatgattcaattagcttccgactcattcatccaaataccgag tcgccaaaattcaaactagactcgttaaatgaatgaatgatgcggtagacaaattggatcattgattctctttga**gcgcgcaaaagagacgtta aa**tctctcttttgtattccaattttcttgattaatctttcctgcacaaaaacatgcttgatccactaagtgacatatatgctgccttcgtatat atagttctggtaaaattaacattttgggtttatctttatttaaggcatcgccatgacccagctttcttgtacaaagtggt*

The *RaichuROP6* constructs were based on Raichu-Rac1 and Raichu-Cdc42 constructs (43-45). In Raichu-Rac1/Cdc42 sensors, the CRIB domain of human PAK1 is cloned in tandem with Rac1 or Cdc42 (interspaced by an appropriate linker) and flanked by two fluorescent protein FRET pairs (Venus and ECFP) (43-45). The CRIB domain interacts specifically with activated Rac1/Cdc42 (i.e. GTP-bound Rac1 or Cdc42), which induces a conformational change in the sensor and enhances the FRET efficiency between the two fluorescent proteins. We used a similar design for the Raichu-ROP6 sensor. Because the CRIB domain of human PAK1 is known to also interact with plants GTP-bound ROPs (18, 23), we used this domain as a generic probe for ROP6-GTP conformation. To clone the *RaichuROP6* constructs, we first ordered a synthetic gene with the following sequence and flanked by *AttB1* and *AttB2* regions and subsequently cloned it into *pDONR221* by BP recombination: *ggggacaagtttgtacaaaaaagcaggcttagaattcggcatggtatccaaaggtgaagaattatttacgggagtcgtgccaatacttgtcgagttggacggtgatgtgaacgggcataaattttcagtaagcggggagggagagggtgacgctacatacggaaaattaactttgaaactaatctgtaccacaggaaaactgcctgtaccgtggcctactctcgtaacaacacttggttacggtttacaatgttttgcacgttaccccgaccatatgaaacaacatgacttcttcaaatctgcgatgcccgagggctatgttcaagaaaggactatattcttcaaggacgacggtaattacaaaactagagctgaagtcaagttcgaaggagacacgctcgtgaataggatcgagttaaagggaatcgacttcaaagaagatggaaacattctgggacacaaattggagtacaactataattcccataacgtttacatcaccgccgacaaacaaaagaacgggataaaggcaaacttcaagattaggcacaatattgaggatgggggagtccagttagccgatcactaccagcaaaatactcccataggtgacgggcccgtgctgcttcccgataatcattatctatcataccagtctgcgcttagcaaagaccctaacgagaagagagatcacatggtcttgttggaattcgtaacagctgccctcgagaaagagaaagagcggccagagatttctctcccttcagattttgaacacacaattcatgtcggttttgatgctgtcacaggggagtttacgggaatgccagagcagtgggcccgcttgcttcagacatcaaatatcactaagtcggagcagaagaaaaacccgcaggctgttctggatgtgttggagttttacaactcgaagaagacatccaacagccagaaatacatgagctttacagataagtcagcttccggaggtggaaccggtggtggaggtaccatgagtgcttcaaggtttatcaagtgtgtcactgtcggcgacggtgctgttggaaagacttgtcttctcatctcctacactagcaacactttccccacggattatgtgccaactgtgttcgataatttcagtgccaatgtgattgttgatggcaacactatcaacttgggattgtgggatactgcagggcaagaggactacaatagactaagacctttgagctatcgcggtgcagatgtcttcttacttgcattctcacttgtcagcaaagctagctatgaaaatgtttctaaaaagtgggttcctgaactgagacattatgctcctggtgttcccatcatcctcgttggaacaaagcttgatcttcgagatgataagcaattctttgccgagcaccctggtgctgtgcctatctctaccgctcagggtgaagaactaaagaagctgattggggcgcctgcttatatcgaatgcagtgcaaaaactcaacagaatgtgaaagcagtgtttgatgcggctatcaaggtcgttcgcggccgcatggtgtccaagggagaagaactttttaccggagtagtacctatacttgtcgagctcgatggggatgtcaatggtcaccgtttctcagtcagcggggaaggcgagggtgacgcgacttacgggaaattaaccctaaaatttatatgcaccacgggaaagctgccagtaccctggcccaccctggtcaccaccctaacctggggcgtccagtgtttctcacgataccctgaccacatgaagcaacacgatttcttcaaatcagctatgcccgaggggtatgtccaggagagaacgatattttttaaggatgatgggaattacaagactagggctgaggttaagttcgaaggggatacgcttgtgaacaggatagagttaaaaggcattggctttaaggaggatggtaacatccttggccacaaactagagtataactacatcagtcataacgtgtacataacggccgataagcagaagaatggtattaaggcacattttaagataagacacaacatcgaagatggtggggtccaactggctgatcattaccagcaaaatactcctatcggcgacggccctgtattgctaccggacaatcactacctaagtactcaaagcgcgttatccaaggaccctaacgagaagagagatcacatggttctgcttgaatttgtgaccgcagctggaatcacgcttggaatggacgaactgtctagagtcgctatcaaggtcgttctccagccaccaaaaaacaagaagaagaagaagagaaaatctcagaaaggttgttctatactctagggatcctacccagctttcttgtacaaagtggtcccc*

*RaichuROP6/pDONR221* was then used as template for PCR amplification by the *ROP6-CA_F* and *ROP6-CA_R* primers to give *RaichuROP6^CA^/pDONR221*.

*RaichuROP6^CA^/pDONR221* was then used as template for PCR amplification by the 5’phosphorylated *RaichuROP6-7Q_F* (*caacaatctcagaaaggttgttctatactctagggatcctacccagctttcttgtac*) and *RaichuROP6-7Q_R (ctgctgctgctgctggttttttggtggctggagaacgaccttgatagcgactctagac*) primers followed by ligation to give *RaichuROP6^CA-7Q^/pDONR221*.

#### Construction of destination clones (destination vector)

The following gateway entry vectors were previously published: *pTNT-HA-ccdb (30); UBQ10prom/pDONR P4P1R (NASC# N2106315) (46), 2×35sprom/pDONR P4P1R (NASC# N2106316) (46), mCITRINEnoSTOP/pDONR221 (NASC# N2106287) (47);* and *2xPH^EVCT2^/pDONR P2RP3 (28*).

Binary destination vectors for plant transformation were obtained using the multisite LR recombination system (life technologies, http://www.thermofisher.com/) using the *pB7m34GW* (basta resistant) or *pK7m34GW* (Kanamycin resistant) (*48*) as destination vectors, and the donor vectors describe above to give: *pROP6prom∷mCITRINE-ROP6/pB7m34GW, pROP6prom∷mCITRINE-ROP6^3Q^/pB7m34GW, pROP6prom∷mCITRINE-ROP6^7Q^/pB7m34GW, 2×35sprom∷mEOS-ROP6/pB7m34GW, 2×35sprom∷mEOS-ROP6^7Q^/pB7m34GW, 2×35sprom∷mEOS-ROP6^CA^/pB7m34GW, UBQ10prom∷mCITRINE-ROP6^C-term^/pB7m34GW, UBQ10prom∷mCITRINE-ROP6^3Q-C-term^/pB7m34GW, UBQ10prom∷mCITRINE-ROP6^7Q-C-term^/pB7m34GW, 2×35sprom∷mEOS-2xPH^EVCT2^/pB7m34GW, promUBQ10∷PSS1-OX-mCITRINE/pB7m34GW, promUBQ10∷PSS1-AMI1/pK7m34GW, promUBQ10∷PSS1-AMI2/pK7m34GW, prom2×35S∷RaichuROP6/pB7m34GW, prom2×35S∷RaichuROP6^CA^/pB7m34GW, prom2×35S∷RaichuROP6^CA-7Q^/pB7m34GW*. Transgenic lines were obtained as described in (46).

Each transgenic line was used as follow:

***UBQ10prom∷mCitrine-C2^LACT^** (30):* Fig. 1A and B.

***UBQ10prom∷mCitrine-2xPH^EVCT2^** (28):* Fig. S1B and C.

***pss1-3*** (28): Fig. 2A-D, Fig. 4A-C, E and F, Fig. 6C, Fig. S1F, Fig. S2A-F and I, Fig. S3A-C and E, Fig. S5G and H, Fig. S10B-F.

***pss1-4*** (28): Fig. S1F and G, Fig. S10B-F.

***pss1-5*** (28): Fig. S1F and G, Fig. S10B-E. *pss1-3xpPSS1∷PSS1g (pss1-3xPSS1prom∷PSS1genomic*) (28): Fig. 2A, Fig. S2A, Fig. S10C and D.

***GFP-ROP6** (2×35S∷GFP-ROP6*) (34): Fig. 3A-C, Fig. 4A, B and D-F, Fig. S3A-D, Fig. S7D, Fig. S10C and D.

***pss1-3xGFP-ROP6**:* Fig. 2A, Fig. 3A-C, Fig. 4A, C, E and F, Fig. S3A-C and E ***UBQ10prom∷2xmCherry-C2^LACT^x2×35S∷GFP-ROP6**:* Fig. 5C

***GFP-ROP6^CA^** (2×35S∷GFP-ROP6-CA) (34):* Fig. S10C and D.

***PSS1-OX** (UBQprom10∷PSS1-mCitrine) and **PSS1-AMI** (UBQ10prom∷AMI1 and 2):* Fig. 2B and D, Fig. S1D and E, Fig. S2G-I.

***CLC2-GFP** (pCLC2∷CLC2-GFP) (39*) and ***pss1-3xCLC2-GFP**:* Fig. S2E. ***GFP-TUA6** (35S∷GFP-TUA6, NASC #6551*) and ***pss1-3xTUA6-GFP***: Fig. S2F. ***mEos-ROP6** (also called **ROP6-OX** for phenotypic analyses, 2×35prom∷mEos-ROP6*): Fig. 2A, Fig. 3D-G, Fig. 6A-B, Fig. S5B and C, Fig. S6B-C and E, Fig. S8D-F.

***pss1-3xmEos-ROP6** (also called **pss1-3xROP6-OX** for phenotypic analyses*): Fig. 2A, Fig. 6C, Fig. S5G and H, Fig. S6E.

***mEos-ROP6^CA^** (also called **ROP6^CA^** for phenotypic analyses, 2×35S∷mEos-ROP6^CA^):* Fig. 2A and B, Fig. 3E and G, Fig. S5A, Fig. S6A and E.

***mEos-ROP6^7Q^** (also called **ROP6^7Q^-OX** for phenotypic analyses,2×35S∷mEos-ROP6^7Q^):* Fig. 6A-C, Fig. S5E and f, Fig. S8D-F.

**mEos-2xPH^EVCT2^** (2×35S∷mEos-2xPH^EVCT2^): Fig. 5A and B, Fig. S5D.

***PIN2-GFP** (40) (pPIN2∷PIN2-GFP*) and ***pss1-3xPIN2-GFP***: Fig. S2B-D. ***rop6-2xPIN2-GFP***: Fig. S2B-C.

***rop6-2** (22):* Fig. S8B and C, Fig. S10C and D. ***rop6-2xROP6prom∷mCitrine-ROP6***: Fig. S7C and D, Fig. S8A and B. ***rop6-2xROP6prom∷mCitrine-ROP6^3Q^***: Fig. S7C. ***rop6-2xROP6prom∷mCitrine-ROP6^7Q^***: Fig. S7C and Fig. S8A and B. ***UBQ10prom∷mCitrine-ROP6-C-term**:* Fig. S7C and D. ***UBQ10prom∷mCitrine-ROP6^3Q^-C-term**:* Fig. S7C. ***UBQ10prom∷mCitrine-ROP6^7Q^-C-term**:* Fig. S7C.

***PIP2a-mEos** (pUBQ10∷PIP2a-mEOS) (15):* Fig. S8F.

#### Recombinant protein expression and lipid-protein overlay assays

The expression plasmids *pTNT∷HA-ROP6cDNA* and *pTNT∷HA-ROP6^7Q^cDNA* were obtained by LR recombination between *ROP6cDNA/pDONR221, ROP6^7Q^cDNA/pDONR221* entry vectors and the *pTNT-HA-ccdb* destination vector. The expression plasmids *pTNT∷HA-ROP6cDNA* and *pTNT∷HA-ROP6^7Q^cDNA* were used as DNA template for *in vitro* transcription and translation using the TNT^®^ SP6 High-Yield Wheat Germ Protein Expression System (Promega, www.promega.com), following manufacturer’s instructions. 5μl of the total reaction were used to analyze protein expression levels by western-blot using 1:1000 anti-HA (www.boehringer-ingelheim.com) primary antibody and 1:5000 secondary anti-mouse (GE Healthcare Life Sciences, http://www.gelifesciences.com/) antibody. The lipid overlay assays were performed as follow: nitrocellulose membranes containing immobilized purified lipids (PIPstrip P-6001, Echelon Bioscience, http://echelon-inc.com/) were incubated for 60 min in blocking solution (TBST (50 mM Tris, 150 mM NaCl, 0,05% Tween 20, pH 7.6) + 3% BSA). Membranes were then incubated for 2h with 10mL of blocking solution containing 40 μl of in vitro synthesized proteins. After three washing steps using blocking solution, membranes were incubated for 120 min at room temperature with primary antibodies diluted in blocking solution, rinsed three times with blocking solution and incubated for 60 min at room temperature with the secondary antibody also diluted in blocking solution. Antibodies and dilutions are the same as described above.

#### Recombinant protein expression and purification for enzymology assay

The full length ROP6 WT and 7Q coding sequence were inserted in pET28a+ between NdeI and NotI for overexpression in Escherichia coli (DE3) Rosetta pLysS (Novagen, www.merckmillipore.com/). The bacterial culture (2xTY medium, www.sigmaaldrich.com) from overnight preculture was incubated at 37°C. The induction was performed at OD 0.55 with 0.5mM IPTG + 5% glycerol and incubated at 18°C overnight. The cells were pelleted, resuspended in 20mM Tris, 400mM NaCl, 5mM beta-mercaptoethanol and flash frozen. After thawing, the cells were lysed by sonication and centrifuged at 15,000g for 35 minutes at 4°C. The supernatant was applied on a Nickel Immobilized Metal ion Affinity Chromatography at room temperature. The beads were washed with 10 volumes of 20mM Tris, 400mM NaCl, 20mM imidazole and then with 5 volumes of 20mM Tris, 400mM NaCl, 60mM imidazole. The beads were then eluted with 20mM Tris, 150mM NaCl, 0.4M imidazole. The elution fraction was centrifuged at 20,000g for 20 minutes at 4°C before purification by the Size Exclusion Chromatography (SEC) column Superdex 75 16/60 in 20mM Tris, NaCl 150mM pH 8 at 1mL/min. The fractions corresponding to ROP6 both pure and non-aggregated are pooled and concentrated at 50 μM.

#### GTPase assay

The ROP6 proteins used for the assays were never frozen and used less than 24 hours after their purification. GTP hydrolysis was detected using Biomol Green (Enzo lifescience, www.enzolifesciences.com/) following the 1mL sample commercial protocol. The concentrations used for the reactions are 80μM GTP, 10μM ROP6, 10μM BSA in 20mM Tris, NaCl 150mM, 2mM MgCl2 pH 8. The GTPase reactions were incubated at 22°C for 75 minutes. The measures were repeated three time, with independent ROP6 purification and independent phosphate calibration for each set of experiments.

#### qRT-PCR

Total RNA was extracted using the Spectrum Plant Total RNA Kit (Sigma). Total RNAs were digested with Turbo DNA-free DNase I (Ambion) according to the manufacturer’s instructions. RNA was reverse transcribed using the SuperScript VILO cDNA Synthesis Kit (Invitrogen) according to the manufacturer’s protocol. PCR reactions were performed in an optical 396-well plate in theQuantStudio 6 Flex Real-Time PCR System (Applied Biosystems), using FastStart Universal SYBR Green Master (Rox) (Roche), in a final volume of 10 μl, according to the manufacturer’s instructions. The following standard thermal profile was used for all PCR reactions: 95 °C for 10 min, 40 cycles of 95 °C for 10 s, and 60 °C for 30 s. Data were analyzed using the QuantStudio 6 Flex Real-Time PCR System Software (Applied Biosystems; www.thermofisher.com).

Two set of primers were used to amplify *PSS1* cDNA, the first set of primers (black dots on Fig. S1D) *ggcttacaagcctcgcactatc*/*tcaagagctccactagcccaaatg* was located at the *5’* end of *PSS1* cDNA. The second set of primers (grey dots on Fig. S1D) *gcattctgttggctgtcactgg*/*ttctgttgggtacaatccacttcc* was located at the *3’* end of the *PSS1* cDNA. As a reference, the primers *gactacagtccactcaatcactgc/aagagctggaagcacctttccg* were used to amplify the *GAPC1* (At3g04120) cDNA. PCR efficiency (*E*) was estimated from the data obtained from standard curve amplification using the equation *E*=10^−1/slope^. Expression levels are presented as E^-CtPSS1^ / E^-CtGAPC1^.

### LIPID QUANTIFICATION (HPTLC)

#### Lipid extraction

12 days old seedlings (0.1-1g fresh weight) were collected in glass tubes; 2 ml of preheated isopropanol were added and tubes were heated at 70°C for 20 min to inhibit phospholipase D activity. 6 ml of chloroform/methanol 2/1 (v/v) were added and lipid extraction was completed at room temperature. The organic phases were transferred to new glass tubes. Then 1.5 ml of H2O was added to the organic phases and tubes were vortexed and centrifuged at 2000rpm; the organic phases were transferred to new glass tubes, evaporated and the lipids were resuspended in the appropriate volume of chloroform/methanol 2/1, v/v, in order to obtain the same concentration according to the initial seedlings fresh weight.

#### High performance thin layer chromatography (HPTLC)

Lipids were deposited on HPTLC plates (Silica gel 60G F254 glass plates Merck Millipore) together with external pure lipid standards (Avanti lipids). Plates were developed according to (49). Following chromatography, the lipids were charred for densitometry according to (50). Briefly, plates were dipped into a 3% cupric acetate (w/v)-8% orthophosphoric acid (v/v) solution in H2O and heated at 110°C for 30min. Plates were scanned at 366 nm using a CAMAG TLC scanner 3. 6 independent samples were quantified for wild type, *PSS1-AMI1, PSS1-AMI2, PSS1-OX1* and *PSS1-OX2* plants.

#### Western blot

20μl of the total reaction were used to analyze protein expression levels by western-blot using 1:2000 anti-Eos (A010-mEOS2, https://badrilla.com) and 1:10000 anti H3 (www.boehringer-ingelheim.com) primary antibodies incubated overnight at 4°C. 1:5000 secondary anti-rabbit-HRP antibody was applied at room temperature for 60 min (www.thermofisher.com). For revelation, ECL prime was applied for 30 seconds.

### IMAGE QUANTIFICATION

#### Gravitropic response and gravitropic defects

7-8 days old seedlings were subjected to 135° angle for 12 hours. For each genotype analyzed at least 30 roots in three independent experiments were used for quantification. Every 4 hours, plates were scanned with EPSON scanner perfection V300 PHOTO at 800 dpi. Each plate at the different time points were cropped and aligned using “Template Matching and Slice Alignment” plugin on FIJI to obtain a timelapse for 12h. To allow high throughput data analyses, the process has been automatized on a macro. To quantify the average root angle of curvature RootTrace software was used (51). In Fig. 2A, 2B and 6A, statistical analyses were performed on the root bending at 8 hours after reorientation.

The horizontal and vertical growth index were calculated on 10 days old seedlings of WT, *pss1-3*, *pss1-4* and *pss1-5* mutant plants using the “segment line” tool on FIJI. Briefly, to calculate the gravitropic indexes, three length are considered, L, Ly and Lx (52). L is the total length of the roots (from base of hypocotyl to root tip), while Ly and Lx are, respectively, ordinate and abscissa of the root tip (considered from the point of view of the base of hypocotyl). The horizontal growth index (HGI) corresponds to the ratio Lx/L, while the vertical growth index (VGI) corresponds to the ratio Ly/L (52). For WT and *pss1-3* at least 44 roots in three independent experiments were used and one experiment for *pss1-4* and *pss1-5*. In each graph relative to gravitropic response and defect, “n” represents the number of roots used for quantification.

#### Size of FM4-64-stained BFA bodies

BFA body size was quantified on at least 14 roots representing an average of 602 cells in three independent experiments. Threshold was applied, images harboring less than ten BFA bodies were removed from the analysis as well as images issue from misshapen root cells. Cells containing BFA bodies whose length are bigger than width were selected for the analysis. Images were submitted to a NidBlack auto local threshold with a radius of 15. BFA bodies bigger than 5.2 um^2^ with a circularity between 0.25 and 1 were detected using the “Analyse Particules” plugin of FIJI. The average size of BFA bodies was obtain for one root. Per root an average of 38 BFA bodies was detected representing at least 532 BFA bodies quantified per conditions. All this process has been automatized using a macro. In each graph relative to the size of BFA body, “n” represents the number of root used for quantification

#### Number of PIN2-GFP BFA-body per cells

10-12-day old seedlings expressing PIN2-GFP in WT, *rop6-2* and *pss1-3* were used to quantify the number of cells containing PIN2 BFA bodies. At least 6 roots representing 265 cells in one experiment were used for quantification (Fig. S2). For the concomitant treatment BFA and NAA in Fig. S2, for WT and *pss1-3* at least 29 roots representing 1734 cells in three independent experiment (except for *rop6-2* one experiment) were used for quantification. The percentage of cell was obtained counting by eyes the number of PIN2 BFA divided by the number of cell in the root meristematic and elongation zones multiplied by one hundred. In each graph relative to the number of PIN2-GFP BFA-bodies per cell, “n” represents the number of roots used for quantification.

#### Percentage of CLC2-GFP at plasma membrane (PM)

PM-positive CLC2-GFP were counted by eyes on at least 21 roots of 10-12 days of seedlings, representing 698 cells in three independent experiments. The percentage of cells with CLC2 at the PM was obtained counting the number of PM-positive CLC2 divided by the number of cells in the root meristematic zone multiplied by one hundred. In the graph relative to the percentage of CLC2-GFP at the PM, “n” represents the number of roots used for quantification.

#### Microtubules orientation

Microtubule arrays were acquired on 10-12-day old seedlings in the transition zone of root cells. The average orientation was calculated on at least 28 cells of 13 roots in two independent experiments using FibriTool plugin (53) on Fiji. In the graph relative to microtubule orientation, “n” represents the number of cells used for quantification.

#### Number of spots per cell

The number of spots per cells was calculated using SiCESpotDetector.ijm plugin (*54*) (http://www.enslyon.fr/RDP/SiCE/METHODSfiles/SiCE%20SpotDetectorV3.ijm) on 5 to 7day old transgenic plants expressing ROP6, ROP6^3Q^, ROP6^7Q^ full length fused to mCITRINE exppressed by its own promoter or the ROP6 C-terminal wild type, 3Q and 7Q fused to mCITRINE expressed under the *UBQ10prom*. The quantification was made on at least 12 roots representing an average of 516 cells. In the graph relative to the number of spots per cell, “n” represents the number of roots used for quantification.

#### Fluorescent intensity at plasma membrane (PM)

5-7-day old transgenic lines was used to quantified the plasma membrane intensity according to the integrated density. The integrated density at the plasma membrane was average from 60 cells taken from three independent roots using a three-pixel line width and the “Measure” tool in Fiji. In graphs relative to the fluorescence intensity at the PM, “n” represents the number of cells used for quantification.

#### SptPALM data quantifications and representations

For each movie corresponding to one condition/genotype, the natural logarithm of apparent diffusion coefficient (log(D)) per molecule was extracted from MTT data (42) by CBS sptPALM analyser (Martinière et al., unpublished). For sptPALM analyses, only tracks with at least five successive points were selected and the rest of the molecules were discarded from the analysis. We pooled values from each condition in order to get the log(D) distribution using R software (as histogram shown on left panel of Fig. S5, number of bins set to 80). A first partition P was done; the set of molecules which present a log(D) value below −1.75 refers to the immobile group, whereas values above this threshold denote the mobile group. To obtain the percentage of molecules in the immobile group, we divided the number of tracks below −1.75 by the total track number multiplied by one hundred. Then, each condition was analyzed using an R script to automatize the analysis. To statistically test the existence of these two behaviors (mobile and immobile), we analyzed the log(D) distribution per condition using the mclust R package. For each condition, the percentages of data in each of the 2 groups defined by the partition P with the absolute log(D) value (−1.75) are first reported. We then fitted two distinct two-component Gaussian mixture models, M0 and M1, using the mclust∷Mclust R function. M0 is a model where σ1 ≠ σ2 and M1 is a model where σ1 = σ2. For each model, every molecule is assigned to one of the 2 Gaussian components Gi (i Î [1, 2]) according to a MAP (Maximum A Posteriori)-based classification. M1 model was used since it provided more representative results despite of a lower Bayesian information criterion values. The percentage of values assigned to each of the two Gaussian components is reported as well as its 95% confidence interval obtained with a non-parametric bootstrap method, using the mclust∷MclustBootstrap R function with default arguments. The classification rate was calculated as the percentage of MAP-classified values concordant with the partition P. Based on the estimation of the model M1, we plotted the density of the Gaussian mixture according to the kernel density estimation (called “occurrence” for more clarity) and used for representation (as Fig. S5, right panel). At least 14 acquisitions from three independent experiments were used, representing 12,824 tracks. In each graph relative to the percentage of immobile ROP6, “n” represents the number of acquisitions used for quantification. Note that one acquisition may represent more than 1 cell but that different acquisitions are always from different cells.

#### LivePALM, Voronoi tessellation analyses and representation

For each movie corresponding to one condition/genotype, the object localization was extracted from MTT data (42) by CBS sptPALM analyser (Martiniere et al., unpublished). SR-tesseler software was used to reconstruct live PALM images and determine protein local density (37). Multiple detections of single object were corrected as previously described by Levet et al. (37). Local density was extracted from Voronoï segmentation of the image as shown in Fig. 3F, 5A and Fig. S6. We pooled values from each condition in order to get the log(local density) distribution using R software (as histogram in Fig. S6, left panel, number of bins set to 80). A first partition P was done; the set of molecules which present a log(local density) value below 3.5 refers to the immobile group, whereas values above this threshold denote the mobile group. To obtain the percentage of molecules in the immobile group, we divided the number of molecules below 3.5 by the total molecules number multiplied by one hundred. Then, each condition was analyzed using an R script to automatize the analysis. To statistically test the existence of these two behaviors (mobile and immobile), we analyzed the local density distribution per condition using the mclust R package. For each condition, the percentages of data in each of the 2 groups defined by the partition P with the absolute log(local density) value (3.5) are first reported. We then fitted two distinct two-component Gaussian mixture models, M0 and M1, using the mclust∷Mclust R function. M0 is a model where σ1 ≠ σ2 and M1 is a model where σ1 = σ2. For each model, every molecule is assigned to one of the 2 Gaussian components Gi (i Î [1, 2]) according to a MAP (Maximum A Posteriori)-based classification. M1 model was used since it provided more representative results despite of a lower Bayesian information criterion values. The percentage of values assigned to each of the two Gaussian components is reported as well as its 95% confidence interval obtained with a non-parametric bootstrap method, using the mclust∷MclustBootstrap R function with default arguments. The classification rate was calculated as the percentage of MAP-classified values concordant with the partition P. Based on the estimation of the model M1, we plotted the density of the Gaussian mixture according to the kernel density estimation (called “occurrence” for more clarity) and used for representation (as Fig. S6, right panel). At least 24 cells from three independent experiments were used representing 1,319,164 molecules. In each graph relative to the fraction of clustered ROP6 in percentage, “n” represents the number of roots used for quantification.

#### Clusters density and size analysis

For each movie corresponding to one condition/genotype, the local density of each molecule was calculated with SR-Tesseler (37). For cluster detection, a threshold of 20 times the average localization density was applied to identify regions with high local density. “Objects” with at least 5 molecules were identified as “clusters” of molecules (as shown in Fig. S6D) and from which the size of each cluster was extracted (as shown in Fig. S6E). n indicates the number of detected clusters.

#### GFP-ROP6 domain density

An ImageJ macro was written to manually define homogenous area in plasma membrane on TIRF images. ROP6 microdomains were then manually selected using the multipoint tool of ImageJ and added to a result table with corresponding computed domain density. Cell position in Arabidopsis root (Meristem of Elongation zone) was systematically registered after ROI selection for further analysis. In each graph relative to GFP-ROP6 domain density, “n” represents the number of region of interest (ROI) used for quantification.

For the quantification of domain recovery after photobleaching (Fig. 4E), the total number of bleached GFP-ROP6 nanodomains were manually counted as well as the number of GFP-ROP6 nanodomains that recovered fluorescence. “n” represents the number of nanodomains that were present in the bleached region of interest.

#### Ratio PM/cytosol quantification

To determine the localization in the basal meristem (BM) and in the elongation zone (EZ) but also the effects of NAA on PS biosensors mCITRINE-C2^LACT^ and mCITRINE-2xPH^EVCT2^ we calculated and analyzed the “Ratio PM/cytosol fluorescence intensity”. This correspond to the ratio between the fluorescence intensity (Mean Grey Value function of Fiji software) measured in two elliptical region of interest (ROIs) from the plasma membrane region (one at the apical/basal PM region and one in the lateral PM region) and two elliptical ROIs in the cytosol. We quantified in 150 cells over three independent replicates (50 cells per replicate) for mock and NAA and two replicated for BM and EZ. This ratio reveals the degree of localization of the fluorescent reporters from the plasma membrane to the cytosol/endosomes, between the mock and perturbed conditions or BM and EZ. In each graph relative to the ratio PM/cyto fluorescence intensity, “n” represents the number of cells used for quantification at least 17 roots were used.

#### Ratiometric FRET measurement

Interaction between ROP6 and PAK-CRIB domains within the raichu sensors was evaluated by FRET calculation from the CFP to YFP. In each field of view, two pictures were recorded. The first corresponds to the CFP channel and give the donor fluorescent (Ex458, Em467-493). Whereas the second one is the FRET channel called YFP^FRET^ (Ex458, Em 538-573). The ratio image, YFP^FRET^/CFP, was calculated with Fiji software. Mean Grey value of each cells present in the field of view was measured independently by drawing specific ROI. At least 10 fields of view out of two independent transformation assays were analyzed. In graph relative to FRET measurements, “n” represents the number cells used for quantification.

#### Pavement cell circularity

Stage 3 leaves of 28 days old plants were used for pavement cell circularity quantification. For image acquisition, adaxial leaf epidermis was printed on tepid agar at 3% poured on a coverslip. 5 days after printing, pavement cell edges were observed on the slip using Zeiss IMAGER M5 AXIO optical microscope with 40x/0.75 Zeiss EZ. plan-NEOFLUAR objective with DIC illumination. At least 79 pavement cells of 5 independent leaves were analyzed with Fiji using the circularity measurement. In the graph relative to the pavement cell circularity, “n” represents the number of pavement cells used for quantification.

#### Root hair phenotype and initiation site ratio

Root hair phenotyping and initiation site were observed on 5 days old seedlings on at least 7 plants representing 336 root hairs for phenotyping and 48 for initiation site in two independent experiments. In order to determine the initiation site ratio, the length of the root hair initiation (l) from the basal side divided by the total length of the trichoblast (L) were measured. For image acquisition, plants were set up between glass and coverslip containing water and observed using Zeiss IMAGER M5 AXIO optical microscope with 40x/0.75 Zeiss EZ.plan-NEOFLUAR objective. In each graph relative to root hair phenotyping and initiation site, “n” represents the number of root hair cells used for quantification.

#### Lateral root density

12 days old seedlings were used for quantification for the complementation assay, plates were scanned with EPSON scanner perfection V300 PHOTO at 800 dpi. A ratio of the number of lateral roots divided by the root length was applied in order to calculate the lateral root density. At least 116 plants were analyzed in two independent experiments. In the graph relative to the lateral root density, “n” represents the number of roots used for quantification.

### STATISTICAL ANALYSES

Each sample were subjected to four different normality tests (Jarque-Bera, Lilliefors, Anderson-Darling and Shapiro-Wilk), sample were considered as a Gaussian distribution when at least one test was significant (p=0.05). Consequently, parametric or non-parametric test were performed. For parametric test, an ANOVA was performed coupled to a Fisher test in order to proceed to pairwise comparison between samples (confidence index, 95%). Statistical analyses between two samples were performed using the Student t-test (p-value=0.05 or 0.10 in the case of gravitropism experiment). For non-parametric test, results were statistically compared using the Kruskal-Wallis bilateral test (p-value=0.05) using XLstat software (http://www.xlstat.com/). Pairwise comparisons between groups were performed according to Steel-Dwass-Critchlow-Fligner procedure (different letters indicate statistical difference between samples) (Hollander and Wolfe, 1999). Statistical analyses between two samples were performed using the non-parametric Wilcoxon-Mann-Whitney test (p-value=0.05).

The following tests were performed for each graph in the main Figures and Supplementary Figures:

#### Non-parametric test

Kruskal Wallis bilateral test combined with a multiple pairwise comparisons:

- Using the Conover-Iman procedure / Two-tailed test, significance level 15%: Fig. 2A
- Using the Steel-Dwass-Critchlow-Fligner, significance level 5%: Fig. 3G, Fig. 4B-C and E, Fig. S2E, Fig. S8C (10%), Fig. S10C.

Wilcoxon-Mann-Whitney /Two-tailed test, significance level 5%: Fig. 1A-B, Fig. 3C, Fig. S1B-C.

#### Parametric test

Student t-test for two independent samples / Two-tailed test, significance level 5%: Fig. 4D. Two ways ANOVA, confidence index 95% combine with an analysis of the differences between the categories:

- Using the Fisher test: Fig. 2B-C, Fig. 3E, Fig. 4A, Fig. 6B-C, Fig S1D, Fig S1E, Fig. S2C-D, F and Hh
- Using the Tukey test: Fig. 2D, Fig. 6A, Fig. S2A (85%), Fig. S7C, Fig. S8B,

